# Sex- and age-specific body condition decline under warmer and drier breeding conditions in European robins

**DOI:** 10.64898/2026.06.07.725434

**Authors:** Mara López-Zuluaga, Carolina Remacha, Ana Bermejo-Bermejo, Emilio Escudero, Javier de la Puente, Javier Pérez-Tris

**Author notes:** Carolina Remacha and Mara López-Zuluaga should be considered joint first author.

## Abstract

1. Local environmental conditions during the breeding season can limit bird populations. Identifying which variables, when, and how they affect key biological traits such as body condition is crucial for understanding long-term population trends under ongoing climate change.
2. We analysed the relationships between environmental variables and body condition during the breeding season in European robins (*Erithacus rubecula*), aiming to uncover links between short-term environmental influences and long-term trends in body condition in the context of local climate change.
3. Using data from a robin population monitored between 2007 and 2021, we applied weather sliding-window analyses to identify periods when temperature, soil moisture, and vegetation productivity best predicted individual body condition. For each variable, we identified critical time windows (CTWs) influencing (1) body condition across the season and (2) individual changes within two weeks. Juveniles and adults were analysed separately, with adult males and females distinguished during pre- and post-fledging periods. We also assessed long-term trends in environmental variables and body condition, and examined how body condition was correlated with apparent survival.
4. Body condition variation across the season was explained by different environmental variables depending on age, sex, and period. Body condition declined with increasing minimum temperatures in adult males and juveniles, and with low soil moisture in adults of both sexes. We did not identify reliable CTWs explaining short-term within-individual changes in body condition. Across 2007–2021, body condition in adult males during the post-fledging period declined with rising minimum temperatures, while fledging dates advanced. Apparent survival was positively associated with body condition only in juvenile robins.
5. Our results reveal multiple seasonal environmental influences that may contribute to short- and long-term declines in body condition in European robins, with effects particularly strong (or most detectable) in adult males. Reduced body condition may have demographic consequences by lowering juvenile survival, although shifts in breeding phenology could mitigate this impact. Overall, these findings highlight how environmental effects on body condition can shape long-term population trends and species vulnerability to climate change.

## INTRODUCTION

There is great concern about how populations, species, and ecosystems may be vulnerable to climate change (Visser & Gienapp, 2019). Birds are directly constrained by environmental variables through the alteration of food availability or physiology, which can determine individual survival and breeding performance (Both, 2010; Pearce-Higgins, 2010). Phenological adjustments (Dunn & Winkler, 2010), abundance or fitness declines (Both et al., 2006), and range shifts (Bateman et al., 2016) are common population responses to changing environment, which are mediated by short-term biological responses of individuals, such as hatching failure or impaired body condition associated with adverse weather conditions. Therefore, knowing the tempo and mode of short-time responses can help us to link individual performance with population vulnerability to climate change (Van De Pol et al., 2016).

The identification of critical periods during which key biological traits are most affected by weather conditions, known as weather critical temporal windows (Van De Pol et al., 2016), may provide valuable information for assessing how bird species respond to environmental shifts. The timing and duration of exposure to adverse environmental conditions may determine the strength of impacts on individual performance, but these effects may be diverse and complex. Short exposure to unfavourable conditions during critical periods can have a stronger influence than extreme conditions during less demanding periods (Nord & Giroud, 2020), and the same environmental variable can have different effects depending on whether exposure is recent or long-term (Kruuk et al., 2015). In addition, individual responses may be non-linear, reaching maximum or minimum phenotypic values under intermediate environmental conditions (McLean et al., 2018; Yom-Tov et al., 2006), and different traits may show variable sensitivity and critical periods of exposure to weather (Foden et al., 2013). For example, in some tropical bird species egg laying may be triggered by small rain events occurred a few weeks earlier (Hidalgo Aranzamendi et al., 2019), while fledging success depends on short-term exposure to rainfall during incubation (Ortega et al., 2022). Additionally, the same weather variable can have opposite effects on different traits, such as precipitation favouring clutch size but reducing fledgling body mass in Barn owls *Tyto alba* (Chausson et al., 2014).

Predictions of species responses under different climate change scenarios may improve when the underlying biological mechanisms are incorporated in the models (Zipkin & Williams, 2025), but most long-term monitoring schemes are focused on numerical trends (Urban et al., 2016). This might be partly explained because obtaining fine-scale fitness measures such as breeding success is unfeasible for most species. However, morphological traits measured during individual monitoring programs can provide an affordable tool to infer the ecological processes underlying the observed patterns (Tellería et al., 2013). In birds, body mass corrected for size is widely used as an index of body condition, as it reflects the nutritional status and thermoregulatory capacity of individuals (Brown, 1996). Body condition may be key to explain variance in individual fitness, but it changes over an individual’s lifetime due to environmental influences (Wilson & Nussey, 2010), and changes in body condition at different ages may have different impacts on survival and breeding success (Blums et al., 2005; Magrath, 1991). An individual’s body condition may also change throughout the day due to biotic (predation risk, foraging resources) or abiotic influences (weather changes; Clark, 1979).

Body condition usually varies in relation to precipitation and temperature, but this relationship may change among species. Individuals may lose weight when temperature increases (McLean et al., 2022; Van Buskirk et al., 2010), especially in arid environments (Du Plessis et al., 2012; Gardner et al., 2016). This loss can be related with physiological changes such as increased evaporative water loss, for example through panting behaviour, or hyperthermia (Nilsson et al., 2016). Rainfall may favour body condition when it increases food availability (Du Plessis et al., 2012; McLean et al., 2018). However, the opposite effects of temperature and precipitation are expected if animals have to invest energy in thermoregulation when temperature decreases or are unable to feed during rainfall periods, especially if fasting coincides with critical life-history stages (Marques-Santos & Dingemanse, 2020). Finally, weather influence on body condition may depend on individual age and sex. Juvenile condition is often closely linked to short-term environmental variation, especially if conditions deteriorate during critical developmental stages (Monaghan, 2008), whereas adults may cope with environmental stochasticity by balancing reproductive investment against self-maintenance, with different optimal outcomes for males and females if parental investment depends on sex (Balbontín et al., 2012; Stearns, 1992).

Environmental influences may accumulate into long-term trends that parallel ongoing climate change, such as the decline in body condition of common birds with increasing temperatures in northern Europe (McLean et al., 2022). During the last half century, global temperature has risen by around 0.2 °C per decade, with more frequent and more intense extreme heat events (Robinson et al., 2021). But climate changes are not globally homogeneous (Arnell et al., 2019), which leads to population-specific trends of change in body condition (Bailey et al., 2022). Models predict a northward shift in the range of some species under expected climate change (Huntley et al., 2008), suggesting that Mediterranean populations of Palearctic bird species whose southern range boundary occurs in this region could be highly vulnerable. However, few studies have monitored phenotypic trends in the Mediterranean, and how local climate change is paralleled by body condition trends is poorly known.

We analysed data from a Mediterranean population of the European robin *Erithacus rubecula* monitored between 2007 and 2021 at a constant-effort ringing site (CES), where weekly capture sessions were conducted during the breeding season. Our aim was to identify associations between environmental variation and individual body condition, both within and across years, and to investigate the demographic consequences of these associations. Robins may be declining at the edge of their Iberian range (Tellería, 2015), and warming temperatures with more severe summer droughts could be the cause (Jylhä et al., 2010). In the Mediterranean, robins prefer moist forests and show declining body condition across gradients of increasing aridity (Pérez-Tris et al., 2000), reflecting their dependence on ground and undergrowth invertebrates (Kirwan et al., 2025). Therefore, we predicted relationships between body condition and environmental variables that describe a warming and drying habitat for robins: temperature (minimum and maximum), soil moisture, and vegetation productivity. We expected that high temperatures, low productivity and low soil moisture will negatively impact body condition, although we allowed for non-linear responses (for example, excessive humidity during floods may decrease food availability). We analysed juvenile and adult body condition separately, in the latter case further distinguishing between males and females, aiming at identifying the time-frame in which environmental variables were most strongly correlated with body condition in each population group, using a critical temporal windows approach (Van De Pol et al., 2016). Scaling up from within-season patterns to long-term trends, we assessed whether local environmental changes were paralleled by changes in body condition of robins during the period 2007–2021. Finally, we investigated the demographic consequences of the observed patterns by analysing the relationship between body condition and apparent survival in robins and explored phenological adjustment as an alternative population response that could buffer environmental impacts on individual performance.

## METHODS

### Study species and field methods

The European robin is a widespread species in the Western Palearctic, which has a continuous distribution on the norther half of the Iberian Peninsula but becomes increasingly restricted to wooded mountain areas further south (Piñeiro, 2022). Mainly monogamous with biparental care, in Mediterranean forests it usually lays two clutches between April and July. Its diet is mainly insectivorous, completed with vegetal food mainly in winter (Kirwan et al., 2025). We monitored a breeding population of robins at a constant-effort ringing station operating from 2007 to 2021 in a mature forest dominated by Pyrenean oak (*Quercus pyrenaica*) with dense shrubby undergrowth in central Spain (La Herrería, 40°34′ N, 4°09′ W, 900 m a.s.l., one of PASER program sites for monitoring common breeding birds in Spain, (Remacha et al., 2021). The area has a continental Mediterranean climate, with mild and wet winters, and warm to hot, dry summers that extend from June to September (Lionello et al., 2006). Annual mean precipitation is approximately 800 mm and mean temperature is 13 °C, with monthly averages ranging from about 5 °C in January to 23 °C in July (Font Tullot, 2000).

Sampling sessions took place approximately each week between April and July (Figure S1). Following constant-effort ringing methodology, birds were collected from nets every hour, or more frequently if weather conditions required, across five capture rounds starting 1 h after sunrise. Individuals were aged as juveniles (birds that hatched that year) or adults (birds in their second year or older) based on plumage colouration and moult (Jenni & Winkler, 2020). Males and female robins have similar appearance, so the presence of a brood patch and the shape of the cloaca was used to sex breeding adults (Svensson, 1992). Gravid females were excluded from the analyses, for which we dropped all body mass data of females with brood patch scored 6 (gravid female) in the field or with body mass exceeding 1.5 SD from the mean, a criterion based on data visualizations informed by observed mass gain due to gravidity of recaptured females and the distribution of females scored as gravid in the field. Birds were weighed with a digital balance (precision 0.01g) and measured tarsus length (precision 0.01 mm).

Individuals were marked with official rings and released unharmed at the site of capture. Recaptured birds were handled following the same procedure, often by different observers on each capture occasion (repeatability of tarsus length estimated as the intraclass correlation coefficient was high: *r*_i_ = 0.86, *P* < 0.001).

### Environmental variables

We obtained remote-sensing data of land surface temperature and soil moisture from the ERA5-Land database (Copernicus Climate Change Service, 2019; Muñoz-Sabater et al., 2021), with temporal resolution = 1 h and spatial resolution = 9 km. To measure vegetation productivity, we used the Enhanced Vegetation Index (EVI; Lobell et al., 2010) available in MOD13 using Google Earth Engine, with temporal resolution = 16 days and spatial resolution = 250 m (Didan, 2015). To calculate daily EVI values, we adjusted an additive generalized model (GAM) of EVI as a function of Julian date (1 = January 1, hereafter date) using the R package *mgcv* (Wood, 2017).

### Statistical analyses

We assessed the body condition variation between and within individuals differentiating three population groups: juveniles, adult males and adult females. Body condition was calculated as the residuals of a linear model of body mass against tarsus length and capture round for each population group (Brown, 1996). Within-year repeatability estimates revealed greater variation in the body condition of adults, especially in females (r_female-adult_ = 0.22, *P* = 0.02, n = 61; r_male-adult_ = 0.40, *P* < 0.001, n = 84), than in juveniles (r_juvenile_ = 0.71, *P* < 0.001, n = 114). We also considered two different seasonal periods to account for the different biological investment during the breeding time (Figure S2.a). The first period refers to the pre-fledging season, ranging between the earliest sampling date and the day before the first juvenile was captured each year (minimum range 93–136 Julian date, January 1 = day 1). The second period (post-fledging season) ranges between the date of capture of the first juvenile and the last sampling day each year (maximum range 137–203 Julian date).

Environmental variables were highly correlated with Julian date (0.25 < |r| < 0.81), due to inherent seasonal patterns, which led to collinearity problems when raw environmental variation and date were analysed together (Figure S2a-c). Therefore, environmental variables were season-adjusted prior to analysing their relationships with body condition. We extracted the seasonal component of each variable with the *seasadj* function from the forecast package (Hyndman et al., 2024), thereby decoupling the seasonal change in body condition due to breeding stage (in adults) or post-natal performance (in juveniles) from the influence of the environmental variable itself. High season-adjusted weather deviations are expected to negatively impact breeding birds (Cohen et al., 2020).

#### Weather sliding windows for seasonal trends

We used the *climwin* package to identify the range of dates before capture time when each season-adjusted environmental variable was most strongly correlated to body condition (critical time windows, hereafter CTW; Bailey & Van De Pol, 2016). In each analysis, we fitted models considering all possible CTW during a given time frame and compared them using the Akaike information criterion adjusted for small sample size (AICc). Candidate models included the mean value of the environmental variable during each CTW as the environmental predictor, either with linear or quadratic effect to allow for different-shaped environmental influences on body condition, added to a baseline model with covariates relevant to each analysis. We looked for CTW relative to the date of capture because body condition varies seasonally (Clark, 1979), Figure S2.a). To avoid false positives, correct *p*-values (*P*_ΔAICc_) were computed comparing the ΔAICc of best models (those with ΔAICc < −2, where ΔAICc for model *i* = AICc*_i_* – AICc*_baseline,_* preserving the original order of model comparison in *climwin*) with an expected distribution of ΔAICc values obtained from 100 random models with no relationship between weather and body condition. We searched best CTW either for (i) between-individual differences in body condition at the time of capture, and (ii) short-term within-individual changes in body condition, considering in each case the time frame most appropriate for each group of birds.

In the between-individual approach, the baseline model had the following formula:

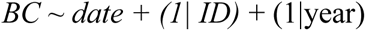

where BC is the body condition of the individual at each capture occasion, date is the Julian date of capture (a fixed covariate that controls for seasonal variation in body condition), and individual identity (ID, as various birds where recaptured within and between years), and year are random effects. For juveniles, we searched CTW during a short time frame of 15 days prior to capture, to cover the nestling period of earliest fledglings. For adults, we explored a longer time frame (30 days prior to capture) to allow longer-term environmental influences without extending into harsh winter conditions for earliest captured individuals.

In the within-individual approach, we analysed the difference in body condition between all pairs of captures of the same bird occurred within a period of up to 14 days, and searched CTW during the 14 days prior to the last capture to assess the influence of environmental events occurred near that date. We used two baseline models:

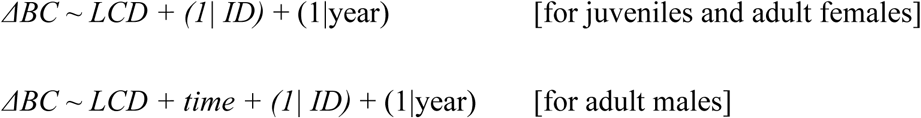

where ΔBC is the individual increment in body condition (positive or negative), LCD is the last capture date (Julian), and time is the number of days elapsed between captures (random effects are the same as in between-individual models). The time effect was considered only in adult males because it was not significant in adult females or juveniles (Text S1, Table S1, males *p* = 0.04, all other *p* > 0.71, Figure S3). For adults, we did not consider different breeding periods due to limited sample size.

When CTW were supported for different environmental variables, we analysed pairwise correlations between these variables to investigate independent influences. If the variables were highly correlated (r > 0.6), we used the one in the model with lowest AICc as representative of a shared influence. If the correlation was lower, we tested each environmental variable with a new baseline model to assess the persistence of its effect controlling for the other significant environmental influences (Hidalgo Aranzamendi et al., 2019; Van De Pol et al., 2016). Finally, when both linear and quadratic effects of one variable were within ΔAIC < −2, we retained the simple linear relationship unless log-likelihood ratio tests (LRT) deemed the quadratic effect to be more reliable.

#### Long-term trends

Once independent short-term influences on body condition had been identified for one or more environmental variables, we investigated whether the variable involved was linked to long-term changes in body condition (Bailey & Van De Pol, 2016). To this end, we analysed the temporal trend of both, body condition and environmental variables, fitting generalized additive mixed models with autoregressive moving average (GAMMs with ARMA) as implemented in the mgcv R package (Ives et al., 2010; Wood, 2017), using Gaussian distribution and restricted maximum likelihood estimation. We selected the best temporal autocorrelation order (AR) comparing models with LRT, increasing AR sequentially until a higher order did not improve model fit. We included the year as a grouping factor in the autocorrelation structure.

For long-term body condition models, we first assessed, in each period, the existence of a linear temporal trend during the period 2007–2021, fitting a “baseline temporal model” with continuous effects of year and date of capture (Julian), and individual identity as a random factor. Then, we fitted a “full environmental model” adding to the baseline temporal model the environmental variables that had an independent influence according to our sliding-window analyses as covariates. This strategy allowed us to assess (1) the existence of a trend of change in body condition and (2) the independent contribution of environmental variables to that trend, which is important if not only body condition, but also the environmental variables that might drive its long-term variation, vary across years. Continuous predictors were z-standardized to estimate comparable effect sizes. Selected quadratic relationships between environmental variables and body condition were included as smooth terms with k = 3 and fx = TRUE. Collinearity was assessed through concurvity test, with values close to 1 meaning lack of identifiability.

For long-term environmental models, we included year (continuous variable) as smooth term. When a significant smooth effect of year was detected, we refitted the model to assess the significance of a linear temporal trend. Given that our models aimed at uncovering climatic trends (between years) irrespective of breeding dates each year, we divided the data in two time-intervals separated by the earliest Julian date of capture of a juvenile any year, hereafter *early* and *late* season (not to be confused with the year-specific pre- and post-fledging periods used in our sliding-window analyses). EVI models with AR > 1 had convergence problems, but models refitted with annual mean EVI values selected AR = 1 and results were similar.

Finally, we assessed whether robins shifted their breeding phenology during the study period, as this could compensate the possible influence on body condition of changing environmental variables. We used linear models of Julian date of capture of the earliest juveniles as a surrogate of fledging date, considering the first two dates with young captured each year to buffer the influence of anecdotal early fledging events. We excluded years 2010, 2015, 2016 and 2018 (with a sampling gap before one of the two dates relevant to assess fledging phenology) to ensure equal sampling effort all years.

#### Survival models

We analysed if body condition predicts between-year apparent survival of robins applying Cormack-Jolly-Seber (CJS) models suitable for long-term capture-mark-recapture (CMR) data of open populations implemented in the R package *RMark* (Laake, 2013). We analysed birds marked initially as juveniles or adults separately because (1) females and males might have different apparent survival or capture probability (Liker & Székely, 2005) but we only know the sex of adults, and (2) body condition might influence survival differently at each age. We selected the best models (ΔAIC _c_ < 2; Burnham & Anderson, 2002) of apparent survival (Φ) between those combining body condition, time as a continuous variable (*T* to assess survival linear trends among years) and, in models with adults, age (second year or older), sex and the interaction between body condition and sex, thereby testing if the influence of body condition on survival differed between males and females. Adult age class was included in the models as a variable with two categories: second year or older (the two adult age classes that can be distinguished in the field). Nevertheless, if adult age did not significantly predict survival, we refitted the model excluding this variable and including adult birds that could not be accurately aged (n = 51). For capture probability (*p*) we included time as a factor (*t* to assess variation between years), within-year sampling effort and sex (only for adults). We first determined the recapture structure in the saturated survival model that best fitted the data. Then, we defined the best apparent survival structure including only the recapture predictors previously selected in reliable models, i.e. those with ΔAIC < 2 if they excluded the constant model. We included time intervals between years, as we had a sampling gap in 2020 due to the COVID-19 pandemic. To estimate *p* in the design matrix, we included yearly sampling effort as the number of sampling days each year (range 10–15 days, Figure S1). We assessed goodness of fit with the function U-CARE of the R package *R2ucare* (Gimenez et al., 2018).

## RESULTS

### Weather sliding windows for population seasonal trends

In adult males, body condition decreased along the season, but this trend was only evident during the pre-fledging period (estimate ± SE = −0.29 ± 0.03, LRT = 57.9, *p <* 0.001; post-fledging period: LRT = 2.66, *p* = 0.10, Figure S2a). Variation in male body condition during the pre-fledging period was best explained by minimum temperature 13 to 10 days before capture (ΔAIC = −21.91, *p_AIC_* < 0.001, Table S2, Fig S4). Male body condition decreased non-linearly with increasing temperature in this time window until it stabilised (Figure 1). During the post-fledging period, we found a linear positive relationship between male body condition and mean soil moisture 23 to 1 days before capture (ΔAIC = −6.23, *p_AIC_* < 0.001, Figure 1, Table S3, Fig S5).

**Figure 1.**
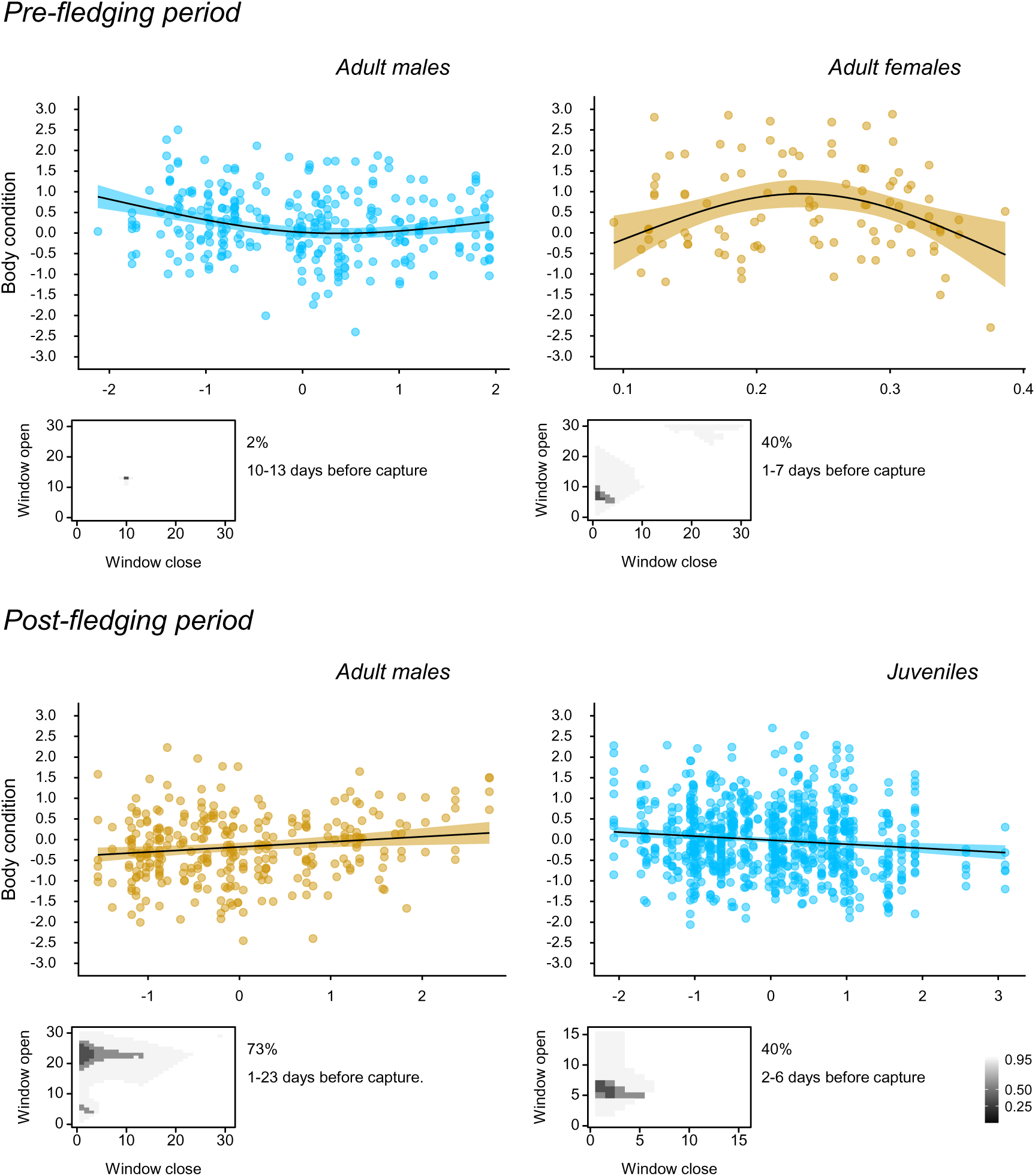
Significant relationships between robin body condition and environmental variables (blue for minimum temperature, ochre for soil moisture) recorded during most reliable critical time windows (CTW) selected by sliding-window analyses in the best model according to ΔAIC. Lines and 95% confidence intervals represent the predictions of generalized additive mixed models (GAMM) with year and Julian date as covariates. The small figure below each pattern shows the time window in the best model and its statistical support measured by the percentage of models that comprise the top 95% of cumulative Akaike weights (which increases when no single CTW stands out over the alternatives, usually due to autocorrelation between candidate models).

In adult females, body condition decreased over the season (pre-fledging period: LRT = 4.46, *p* = 0.03, estimate ± SE = −0.20 ± 0.09; post-fledging period: LRT = 6.82, *p <* 0.01, estimate ± SE = −0.20 ± 0.08; Figure S2a). Variation in female body condition during the pre-fledging period was best explained by soil moisture 7 to 1 days before capture, according to a positive quadratic relationship (ΔAIC = −8.77, *p_AIC_* = 0.01, Table S4, Figure S6) where body condition attained maximum values at intermediate soil moistures (Figure 1). We also found weaker support (ΔAIC = −2.54, *p_AIC_* = 0.01, Table S4) for an increase in body condition with increased EVI the day before capture. We did not detect any environmental influence on body condition in adult females during the post-fledging period (all *p_AIC_ ≥* 0.12, Table S5).

In juveniles, body condition increased along the season (estimate ± SE = 0.009 ± 0.002, LRT = 17.66, *p* < 0.001, Figure S2a). Variation in juvenile body condition was best explained by minimum temperature 6 to 2 days before capture, fitting to a negative linear relationship (ΔAIC = −8.37, *p_AIC_* =0.01, Figure 1, Table S6, Figure S7).

### Weather sliding windows for individual short-term changes

Short-term recaptures (within two weeks) accounted for 44.4%, 30.7% and 77.5% of all captures in adult females, adult males and juveniles, respectively. Individual body mass change between short-term recaptures was 4.27 ± 3.6% (mean of absolute values ± sd) of the original body mass. Within-individual change in body condition was not correlated to any environmental variable during the previous 14 days (all *p* > 0.08, Tables S7-S9), nor was it influenced by the date of the last capture (all *p* > 0.11, for adult males, Text S1, Table S1).

### Long-term trends in body condition

During 2007–2021, body condition decreased in adult males during the post-fledging period (t = −3.16, *p* = 0.002, Figure 2), although this relation was weaker when environmental variables found to be influential in our sliding-window analyses were included in the model (t = −2.16, *p* = 0.03, Table S10). We did not observe long-term trends of change in body condition in adult females or juveniles, nor did we find trends for within-individual short-term change in body condition (all *p* values > 0.07, Table S10).

**Figure 2.**
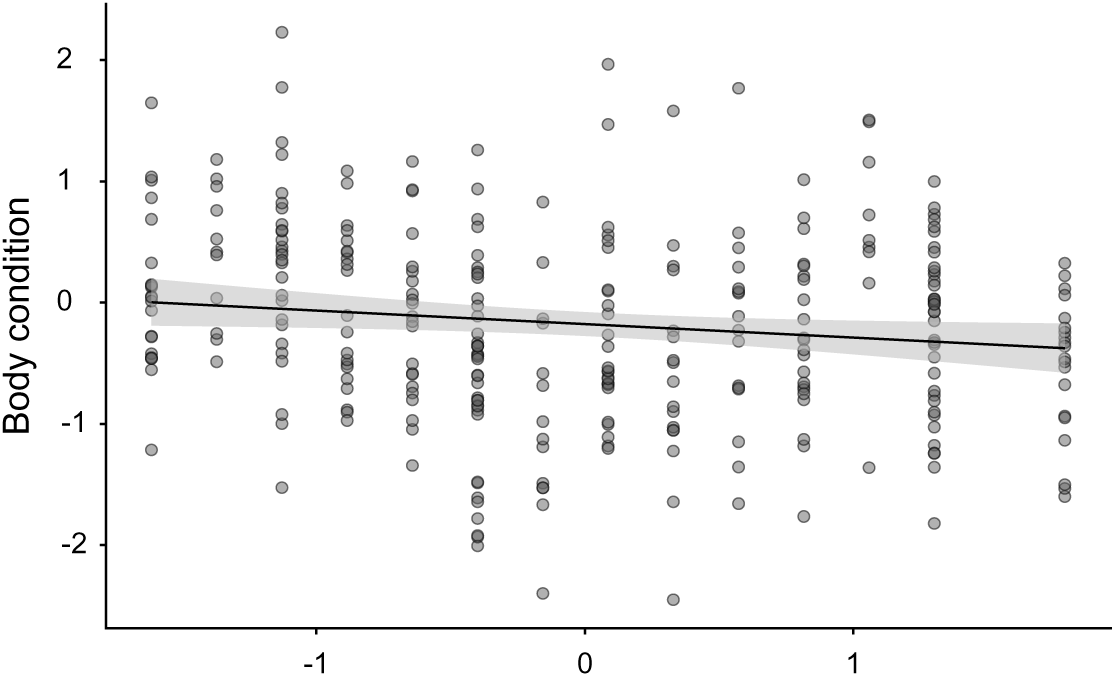
Temporal trend during 2007–2021 of body condition in adult male robins during the post-fledging period. Lines and 95% confidence intervals represent body condition predictions of year (standardized) in a generalized additive mixed model (GAMM) with covariates Julian date and mean soil moisture recorded during the critical time window of 23 days before capture.

### Long-term environmental and phenological trends

Minimum temperature in the post-fledging period increased between 2007 and 2021 (edf = 1.0, F = 4.61, *p* = 0.03, AR = 1). We did not find trends for other variable-period combinations (all *p* > 0.11, Figure S8). The Julian date of capture of the first two juveniles advanced between 2007 and 2021 (F_1,18_ = 4.83; *p* = 0.04, Figure 3a).

**Figure 3.**
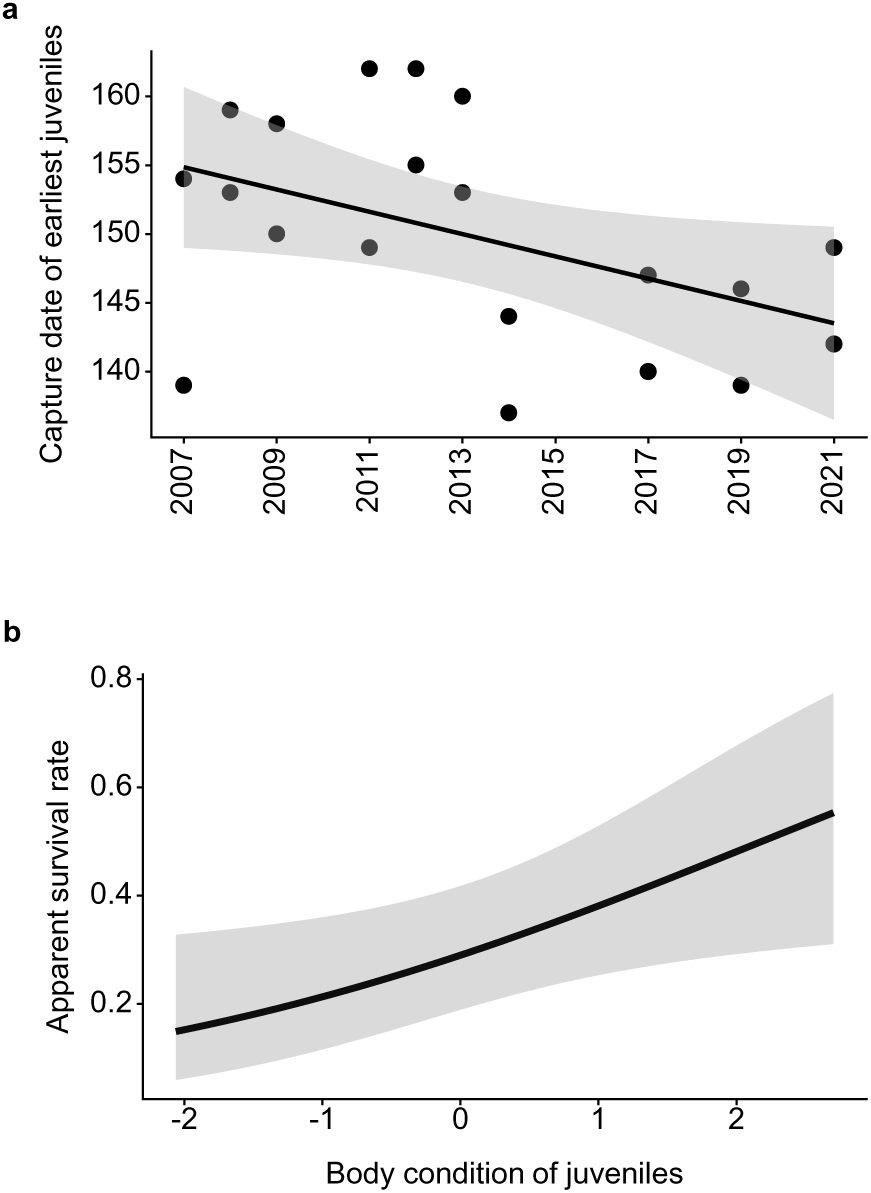
(a) Temporal trend in capture date (Julian date, January 1 = 1) of the two earliest juvenile robins over the period 2007–2021. (b) Relationship between body condition of juvenile robins and apparent survival rate in the best capture-mark-recapture model (Φ _body condition_ + p _intercept_).

### Survival analyses

The percentage of recaptures between years was 4.42% for juveniles and 11.40% for adults. We analysed 1243 capture-recapture histories along the 14 study years (n = 717 juveniles and 526 adults). In adults, survival rates did not significantly vary between second year and older individuals (LRT = 0.10, *p* = 0.75), therefore adults of unknown age were included in our models. Body condition did not determine adult survival between years, but males had higher (estimate ± SE = 35.07 ± 3.80%) apparent survival than females (estimate ± SE = 24.44 ± 4.34%, Table S11). In juveniles, body condition was positively correlated with apparent survival (β = 0.41 ± 0.18, Φ_mean_ ± SE = 28.84% ± 5.90, and recapture probability was not explained by any variable, Figure 3b, Table S11).

## DISCUSSION

Dryer and warmer conditions were associated with lower body condition in robins during the breeding season, with varying effects of each environmental variable in each age, sex and period of the season. Body condition was negatively correlated to minimum temperature in adult males (only during the pre-fledging period) and in juveniles. Soil moisture was positively correlated to body condition in adults, although during different periods in each sex, and with females attaining maximum body condition at intermediate moisture values. General productivity (EVI) had a positive influence on body condition in adult females. In the long term, adult males showed decreasing body condition during the post-fledging period between 2007 and 2021, as minimum temperature increased, although adult apparent survival did not show similar trends and was not predicted by body condition. However, body condition predicted apparent survival in juveniles, which advanced fledging dates between 2007 and 2021. To sum up, our data-driven approach, based on statistical exploration rather than arbitrary decisions, allowed us to uncover the weather signals that best predict body condition in robins (Van De Pol et al., 2016), although such signals were mixed for some environmental variables with a wide range of critical time windows of potential influence. Perhaps for this reason, we did not find any environmental variable related with short-term changes (within two weeks) in individual body condition, although the small sample size for this analysis could have also limited this analysis.

Our results align with the observed abundance distribution of robins in the Iberian Peninsula, where populations face stronger limitation in the warmest and dryest Mediterranean habitats (Tellería & Santos, 1994). An impaired body condition could link environmental variation with robin population limitation for different reasons. First, food availability may become limiting in warming and drying habitats for species that rely on arthropods and earthworms (Martay & Pearce-Higgins, 2020; Pearce-Higgins, 2010). In addition, excessive warming can promote body mass loss in species most sensitive to heat (Du Plessis et al., 2012), either because birds reduce foraging behaviour under high thermal loads (L. Clark, 1987) or because they increase evaporative water loss, which may be especially problematic in small passerines (Albright et al., 2017). Mediterranean robin populations, located at the warmest and driest edge of the species’ range (Pérez-Tris et al., 2000), could be especially vulnerable to the effects of increased temperature and aridity (Jiguet et al., 2010; Pearce-Higgins et al., 2015).

The observed decline in male body condition as local temperatures increased between 2007 and 2021 is consistent with previous studies analysing long-term changes in avian body condition (Campo-Celada et al., 2022; McLean et al., 2022). Besides, we found a negative relationship between temperature and male body condition during the pre-fledging period. Taken together, these results support the idea that climate warming underlies the long-term decline in body condition, although soil moisture could play an additional role in this population of robins. Soil moisture rather than temperature explained variation in male body condition during the post-fledging period, when the summer drought intensifies in Mediterranean habitats (Essa et al., 2023). However, soil moisture did not show the temporal trends observed for temperatures making it less straightforward to establish a link with long-term body condition deterioration. Anyway, the decline in body condition through the years remained detectable after controlling for soil humidity, which suggests that other, not controlled factors also underlie the observed pattern (McLean et al., 2022).

Impaired environmental conditions may impact on robin body condition by reducing food availability (Pearce-Higgins, 2010). In breeding adults, food shortage may further reduce body condition if birds increase food searching effort to secure breeding success (Stearns, 1992). Remarkably, we found a stronger relationship between body condition and environmental variation in male than in female robins. European robins are highly monogamous (Karpińska-Zegarek et al., 2025), with males contributing substantially to mate feeding and parental care, to the point that females often reduce nest provisioning of the first broods to prepare for a second clutch (East, 1981). In addition, males showed lower body condition than females (Figure S2.a), which could make them more vulnerable to environmental influences. In Mediterranean areas, this vulnerability of male robins to habitat deterioration may be particularly important during the post-fledging period, when greater parental load coincides with an intensified summer drought (Essa et al., 2023). However, that we did not observe similar effects in females does not necessarily mean that they are less vulnerable to environmental deterioration. Female robins showed the seasonal decline in body condition typical of passerines, which has been attributed to the thermoregulation cost associated with incubation and brooding (the physiological stress hypothesis; Merila & Wiggins, 1997) or to an adaptive loss of body mass to improve escape performance during predator attacks (the adaptation for flight hypothesis; Forrester & Martin, 2025). Regardless of the cause, females show greater variation in body mass than males across breeding stages. The impossibility of controlling the breeding stage of individuals in our study, along with reduced sample size (females are less likely trapped in mist nets during incubation and brooding periods, and those who were gravid were excluded analyses), may have contributed to lower the detectability of environmental effects in female robins.

While adult body condition decreased seasonally in accordance with the cost of reproduction (Askenmo, 1977), juvenile condition followed the opposite pattern. This result was likely mediated by natural selection favouring heavier individuals, since juvenile body condition was positively related with apparent survival both between years and within years, where recaptured juveniles had better body condition (Text S2, mean ± SD = 0.19 ± 0.96) than not recaptured ones (−0.03 ± 0.81). We found evidence that juvenile robins also gained weight over the post-fledging period, although this mass gain was very small and largely explained by heavy birds recaptured late in the season (see Text S1 for full details).

Body condition in juvenile robins did not show long-term trends of change parallel to increasing temperatures, although minimum temperature predicted juvenile body condition within seasons and juvenile condition predicted apparent juvenile survival. This apparent contradiction could be explained if earlier fledging allowed juveniles to complete development before the summer drought intensifies, thereby compensating the impact of climate warming over the years. Common songbirds, including European robins, are fledging earlier in warmer springs and at warmer sites (Cuchot et al., 2025), a phenological shift that may mitigate climate change impacts (McLean et al., 2022). In fact, bird populations that do not adjust phenology to warming conditions may be more prone to decline (Møller et al., 2008).

The Mediterranean region is considered a climate change hot-spot, where temperature and aridity are expected to continue increasing in the future (Giorgi, 2006). Although we did not find evidence that the observed long-term decline in body condition of adult males translates into reduced survival, we do not know how an intensified drought foreseen in most reliable future scenarios (Essa et al., 2023) may impact on robin populations. Our data suggest that advanced breeding phenology may mitigate climate change impacts to some extent, although more reliable data of fledging dates are required to confirm this trend. In fact, the difficulty to determine the life-cycle stage of individuals is a key limitation of monitoring programs based on bird ringing, compared with more precise methods such as nest-box monitoring. This limitation may have contributed to reduce the detectability of environmental effects in female robins, which show more variable body mass due to clutch formation, egg laying or parental effort, along with their smaller sample size also inherent to the sampling method (females are less likely trapped in mist nets during incubation and brooding periods). However, leveraging the availability of powerful statistical tools (such as time window analysis), we could uncover associations between environmental variation, body condition and survival in robins, which proves the value of standardised ringing as a method that balances sampling effort and quality of information in population monitoring studies.

## Author contributions

M.L.-Z.: Data curation (equal); Formal analysis (supporting); Investigation (equal); Methodology (supporting); Visualization (equal); Writing – original draft (supporting); Writing – review & editing (supporting)

CR: Conceptualization (equal); Data curation (equal); Supervision (equal); Formal analysis (lead); Investigation (equal); Methodology (equal); Project administration (equal); Visualization (equal); Writing – original draft (lead); Writing – review & editing (equal)

A.B.-B: Investigation (supporting); Data curation (supporting); Writing – review & editing (supporting)

E.E.: Investigation (supporting); Data curation (supporting); Writing – review & editing (supporting)

J.P.: Investigation (supporting); Data curation (supporting); Writing – review & editing (supporting)

J.P.-T.: Conceptualization (equal); Data curation (equal); Supervision (equal); Funding acquisition (lead); Investigation (equal); Methodology (equal); Project administration (equal); Resources (lead); Writing – review & editing (equal)

## Acknowledgements

We are grateful to all volunteers involved along the years in the monitoring of breeding birds at La Herrería Bird Observatory, one of the PASER program CES of SEO/Birdlife coordinated by SEO-Monticola Ringing Group. It would have been unachievable to compile this large database without them. Thanks also to Ana Payo-Payo for reviewing our survival analysis. This study was funded by MICIU/AEI /10.13039/501100011033 (grant PID2020-116121GB-I00 to J.P.-T.)

## Data accessibility

Data and R code will be available in Zenodo Digital Repository.

