## Supplementary materials for "Sex- and age-specific body condition decline under warmer and drier breeding conditions in European robins"

3. Present address: BirdWings, Science & Conservation Consulting, El Escorial, Madrid, Spain

### DESCRIPTION OF SUPPLEMENTARY MATERIALS:

- **Figure S1.** Temporal distribution of sampling sessions at La Herrería ringing station between 2007 and 2021. Circles of different sizes indicate the number of Eurasian robins captured in each session, distinguishing adult females (filled red), adult males (filled blue) and juveniles (open circles). Sampling gaps are indicated with N letters, and sampling sessions without robins with blades. During the 14 years of sampling, 1376 different birds were captured (annual mean ± SD = 107.64 ± 32.11 individuals) although effective sample size was reduced due to occasionally missing morphological data.
- **Figure S2.** Seasonal variation in body condition and environmental variables each year. In body condition plots (a), individual values (dots) represent adult females (filled red), adult males (filled blue) and juveniles (filled grey). Vertical dashed lines represent the date of capture of the earliest juvenile each year, which divides each year in *pre-* and *post-fledging* seasons. Raw (b) and season-adjusted (c) values of the environmental variables used as potential correlates of body condition are shown with different colour fill: minimum temperature in blue, maximum temperature in red, soil moisture in ochre and EVI index in green. The vertical line common to all these plots represents the earliest capture date of a juvenile in any year, which sets the limit between *early* and *late* season all years.
- **Text S1. Seasonal changes in within-individual body condition**
  Table S1. Results of repeated-measures mixed models assessing within-individual variation in body mass in relation to Julian date of the last capture and recapture interval (days elapsed between captures). A random factor (recapture time, REC) nested all alternative combinations of first and second capture of each individual each year within bird identity. Degrees of freedom were calculated using the Satterhwaite's method.

- **Figure S3.** Trajectories of body mass change considering all (a) and short-time (b) individual recapture histories within one year for adult males, adult females and juveniles. The sign of the body condition change in each alternative combination is indicated with colours, blue for declining body mass, red for gain and grey for no change. We considered short-time lapse a maximum difference of 14 days between recaptures.
- **CLIMWIN RESULTS:**
  **Tables S2:S9:**

- Identification of critical time windows (CTW) of season-adjusted environmental variables influencing body condition in adult males (**Table S2, S3, S7**), adult females **(Table S4, S5, S8)**, and juveniles **(Table S6, S9)** in the pre- and post-fledgling period (for adults).
- **Table S2.** Identification of critical time windows (CTW) of season-adjusted environmental variables influencing body condition in adult male robins during the pre-fledging period, ranked according to their plausibility based on ΔAIC relative to the baseline model (ΔAICc for model $i$ = $AICc_i$ - $AICc_{baseline}$). The table shows the best model (the one with the lowest ΔAICc) for linear (lin) and quadratic (quad) relationships between the mean value of the environmental predictor during the CTW and body condition. Quadratic relationships are deemed more reliable than linear relationships only when they significantly improve model fit according to LRT tests. The location in time of the CTW is described by the days before capture date when the window opens and closes. Best models are highlighted in bold when significant based on $p_{\Delta AICc}$, resulting from the comparison of model ΔAICc with an expected distribution of ΔAICc values obtained from 100 random models with no relationship between weather and body condition. The percentage of models falling within the 95% confidence set based on cumulative AIC weights is shown only for best-supported environmental influences when significant. Environmental variables with ΔAIC > −2 relative to the baseline model were not further tested.
- **Table S3.** Identification of critical time windows (CTW) of season-adjusted environmental variables influencing body condition in adult male robins during the post-fledging period, ranked according to their plausibility based on ΔAIC relative to the baseline model (ΔAICc for model $i$ = $AICc_i$ - $AICc_{baseline}$). The table shows the best model (the one with the lowest ΔAICc) for linear (lin) and quadratic (quad) relationships between the mean value of the environmental predictor during the CTW and body condition. Quadratic relationships are deemed more reliable than linear relationships only when they significantly improve model fit according to LRT tests. The location in time of the CTW is described by the days before capture date when the window opens and closes. Best models are highlighted in bold when significant based on $p_{\Delta AICc}$, resulting from the comparison of model ΔAICc with an expected distribution of ΔAICc values obtained from 100 random models with no relationship between weather and body condition. The percentage of models falling within the 95% confidence set based on cumulative AIC weights is shown only for best-supported environmental influences when significant. Environmental variables with ΔAIC > −2 relative to the baseline model were not further tested.

- **Table S4.** Identification of critical time windows (CTW) of season-adjusted environmental variables influencing body condition in adult female robins during the pre-fledging period, ranked according to their plausibility based on ΔAIC relative to the baseline model (ΔAICc for model $i$ = $AICc_i$ - $AICc_{baseline}$). The table shows the best model (the one with the lowest ΔAICc) for linear (lin) and quadratic (quad) relationships between the mean value of the environmental predictor during the CTW and body condition. Quadratic relationships are deemed more reliable than linear relationships only when they significantly improve model fit according to LRT tests. The location in time of the CTW is described by the days before capture date when the window opens and closes. Best models are highlighted in bold when significant based on $p_{\Delta AICc}$, resulting from the comparison of model ΔAICc with an expected distribution of ΔAICc values obtained from 100 random models with no relationship between weather and body condition. The percentage of models falling within the 95% confidence set based on cumulative AIC weights is shown only for best-supported environmental influences when significant. Environmental variables with ΔAIC > −2 relative to the baseline model were not further tested.
- **Table S5.** Identification of critical time windows (CTW) of season-adjusted environmental variables influencing body condition in adult female robins during the post-fledging period, ranked according to their plausibility based on ΔAIC relative to the baseline model (ΔAICc for model $i$ = $AICc_i$ - $AICc_{baseline}$). The table shows the best model (the one with the lowest ΔAICc) for linear (lin) and quadratic (quad) relationships between the mean value of the environmental predictor during the CTW and body condition. Quadratic relationships are deemed more reliable than linear relationships only when they significantly improve model fit according to LRT tests. The location in time of the CTW is described by the days before capture date when the window opens and closes. Best models are highlighted in bold when significant based on $p_{\Delta AICc}$, resulting from the comparison of model ΔAICc with an expected distribution of ΔAICc values obtained from 100 random models with no relationship between weather and body condition. The percentage of models falling within the 95% confidence set based on cumulative AIC weights is shown only for best-supported environmental influences when significant. Environmental variables with ΔAIC > −2 relative to the baseline model were not further tested.
- **Table S6.** Identification of critical time windows (CTW) of season-adjusted environmental variables influencing body condition in juvenile robins, ranked according to their plausibility based on ΔAIC relative to the baseline model (ΔAICc for model $i$ = $AICc_i$ - $AICc_{baseline}$). The table shows the best model (the one with the lowest ΔAICc) for linear (lin) and quadratic (quad) relationships between the mean value of the environmental predictor during the CTW and body condition. Quadratic relationships are deemed more reliable than linear relationships only when they significantly improve model fit according to LRT tests. The location in time of the CTW is described by the days before capture date when the window opens and closes. Best models are highlighted in bold when significant based on $p_{\Delta AICc}$, resulting from the comparison of model ΔAICc with an expected distribution of ΔAICc values obtained from 100 random models with no relationship between weather and body condition. The

percentage of models falling within the 95% confidence set based on cumulative AIC weights is shown only for best-supported environmental influences when significant. Environmental variables with ΔAIC > −2 relative to the baseline model were not further tested.

- **Table S7.** Identification of critical time windows (CTW) of season-adjusted environmental variables influencing within-individual variation in body condition in adult male robins, ranked according to their plausibility based on ΔAIC relative to the baseline model (ΔAICc for model $i = AICc_i - AICc_{baseline}$). The table shows the best model (the one with the lowest ΔAICc) for linear (lin) and quadratic (quad) relationships between the mean value of the environmental predictor during the CTW and body condition. Quadratic relationships are deemed more reliable than linear relationships only when they significantly improve model fit according to LRT tests. The location in time of the CTW is described by the days before capture date when the window opens and closes. Best models are highlighted in bold when significant based on $p_{\Delta AICc}$, resulting from the comparison of model ΔAICc with an expected distribution of ΔAICc values obtained from 100 random models with no relationship between weather and body condition. The percentage of models falling within the 95% confidence set based on cumulative AIC weights is shown only for best-supported environmental influences when significant. Environmental variables with ΔAIC > −2 relative to the baseline model were not further tested.
- **Table S8.** Identification of critical time windows (CTW) of season-adjusted environmental variables influencing within-individual variation in body condition in adult female robins, ranked according to their plausibility based on ΔAIC relative to the baseline model (ΔAICc for model $i = AICc_i - AICc_{baseline}$). The table shows the best model (the one with the lowest ΔAICc) for linear (lin) and quadratic (quad) relationships between the mean value of the environmental predictor during the CTW and body condition. Quadratic relationships are deemed more reliable than linear relationships only when they significantly improve model fit according to LRT tests. The location in time of the CTW is described by the days before capture date when the window opens and closes. Best models are highlighted in bold when significant based on $p_{\Delta AICc}$, resulting from the comparison of model ΔAICc with an expected distribution of ΔAICc values obtained from 100 random models with no relationship between weather and body condition. The percentage of models falling within the 95% confidence set based on cumulative AIC weights is shown only for best-supported environmental influences when significant. Environmental variables with ΔAIC > −2 relative to the baseline model were not further tested.
- **Table S9.** Identification of critical time windows (CTW) of season-adjusted environmental variables influencing within-individual variation in body condition in juvenile robins, ranked according to their plausibility based on ΔAIC relative to the baseline model (ΔAICc for model $i = AICc_i - AICc_{baseline}$). The table shows the best model (the one with the lowest ΔAICc) for linear (lin) and quadratic (quad) relationships between the mean value of the environmental predictor during the CTW and body condition. Quadratic relationships are deemed more reliable than linear relationships only when they significantly improve model fit according to LRT tests. The location in time of the CTW is

described by the days before capture date when the window opens and closes. Best models are highlighted in bold when significant based on $p_{\Delta AICc}$, resulting from the comparison of model ΔAICc with an expected distribution of ΔAICc values obtained from 100 random models with no relationship between weather and body condition. The percentage of models falling within the 95% confidence set based on cumulative AIC weights is shown only for best-supported environmental influences when significant. Environmental variables with ΔAIC > −2 relative to the baseline model were not further tested.

- **Figures S4:S9**
- **Fig S4.** Plots describing the results of the best model (the one with the lowest ΔAICc) for the most reliable environmental influence on the body condition in adult male robins during the pre-fledging period (the quadratic relationship with season-adjusted minimum temperature). The figure shows the outcome of the function plotall() of the *climwin* R package, from top left to bottom right: (a) ΔAICc values of models considering every possible critical time window (CTW) during the 30 days before capture when compared to the baseline model without the climatic variable. (b) Percentage of all models falling within the 95% confidence set based on cumulative AICc weights. Coefficients for every CTW of linear (c) and quadratic (d) relationships between season-adjusted minimum temperature and body condition. (e) ΔAICc value of the best model (dashed line) compared with a distribution of ΔAICc values obtained from 100 randomized datasets with no relationship between weather and body condition. (f) Mean window open and close days of the best models included in the 95% confidence set. (g) Most reliable relationship between body condition (biological response) and minimum temperature (climate variable). The best model supported a CTW spanning from day 13 to 10 before capture.

- **Fig S5.** Plots describing the results of the best model (the one with the lowest ΔAICc) for the most reliable environmental influence on the body condition in adult male robins during the post-fledging period (the linear relationship with season-adjusted soil moisture). The figure shows the outcome of the function plotall() of the *climwin* R package, from top left to bottom right: (a) ΔAICc values of models considering every possible critical time window (CTW) during the 30 days before capture when compared to the baseline model without the climatic variable. (b) Percentage of all models falling within the 95% confidence set based on cumulative AICc weights. Coefficients for every CTW of linear (c) and quadratic (d) relationships between season-adjusted soil moisture and body condition. (e) ΔAICc value of the best model (dashed line) compared with a distribution of ΔAICc values obtained from 100 randomized datasets with no relationship between weather and body condition. (f) Mean window open and close days of the best models included in the 95% confidence set. (g) Most reliable relationship between body condition (biological response) and soil moisture (climate variable). The best model supported a CTW spanning from day 23 to 1 before capture, although average CTW open and close days were 21 and 8, respectively.

  **Fig S6.** Plots describing the results of the best model (the one with the lowest ΔAICc) for the most reliable environmental influence on the body condition in adult female robins during the pre-fledging period (the quadratic relationship with season-adjusted soil moisture). The figure shows the outcome of the function plotall() of the *climwin* R package, from top left to bottom right: (a) ΔAICc values of models considering every possible critical time window (CTW) during the 30 days before capture when compared to the baseline model without the climatic variable. (b) Percentage of all models falling within the 95% confidence set based on cumulative AICc weights. Coefficients for every CTW of linear (c) and quadratic (d) relationships between season-adjusted soil moisture and body condition. (e) ΔAICc value of the best model (dashed line) compared with a distribution of ΔAICc values obtained from 100 randomized datasets with no relationship between weather and body condition. (f) Mean window open and close days of the best models included in the 95% confidence set. (g) Most reliable relationship between body condition (biological response) and soil moisture (climate variable). The best model supported a CTW spanning from day 7 to 1 before capture, although average CTW open and close days were 15 and 5, respectively.

- **Fig S7.** Plots describing the results of the best model (the one with the lowest ΔAICc) for the most reliable environmental influence on the body condition in juvenile robins (the linear relationship with season-adjusted minimum temperature). The figure shows the outcome of the function plotall() of the *climwin* R package, from top left to bottom right: (a) ΔAICc values of models considering every possible critical time window (CTW) during the 30 days before capture when compared to the baseline model without the climatic variable. (b) Percentage of all models falling within the 95% confidence set based on cumulative AICc weights. Coefficients for every CTW of linear (c) and quadratic (d) relationships between season-adjusted minimum temperature and body condition. (e) ΔAICc value of the best model (dashed line) compared with a distribution of ΔAICc values obtained from 100 randomized datasets with no relationship between weather and body condition. (f) Mean window open and close days of the best models included in the 95% confidence set. (g) Most reliable relationship between body condition (biological response) and minimum temperature (climate variable). The best model supported a CTW spanning from day 6 to 2 before capture, although average CTW open and close days were 8 and 2, respectively.
- **Table S10.** Results of the models analysing linear long-term trends of body condition in adult males, adult females and juveniles, without (*basic models*) and with environmental variables selected in weather sliding-window analyses (*full environmental models*). Data were fitted with generalized mixed models (*mgcv* package; (Wood, 2017) and included the linear effect of year and Julian date of capture as a basic model. Non-linear patterns were tested (F tests) only when quadratic effects (edf = 2) of environmental variables were deemed most reliable. Year, Julian date and environmental variables were previously standardized. Different models were fit for adults using data from the pre-fledging (PRE) and post-fledging periods (POST). The sample size of each model is indicated (n), and dbc stands for days before capture of critical time windows for environmental influences.
- **Figure S8.** Long-term trends of season-adjusted (sadj) environmental variables (maximum temperature, minimum temperature, soil moisture and EVI) during the *early* (red) and *late* (ochre) breeding seasons across the period 2007– 2021.

- **Table S11**. Best Cormack-Jolly-Seber models (those with ΔAICc < 2) of variation in apparent survival (Φ) and probability of recapture (p) of adult and juvenile robins. We also show the best model that did not include body condition (the variable of interest) and the null model ($\Phi_{\text{intercept}} + p_{\text{intercept}}$). The AIC value, number of parameters (*k*) and Akaike weight are shown for each model.
- **Text S2. Comparison of body condition between birds of each population group that were recaptured or not**

**Figure S1.**

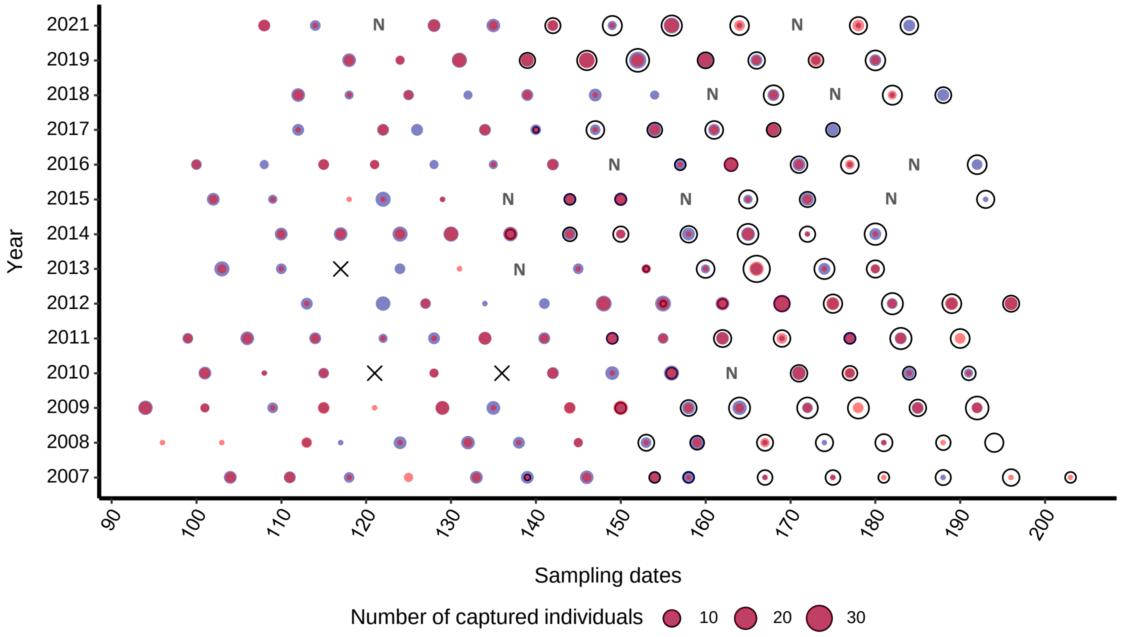


**Figure S1.** Temporal distribution of sampling sessions at La Herrería ringing station between 2007 and 2021. Circles of different sizes indicate the number of Eurasian robins captured in each session, distinguishing adult females (filled red), adult males (filled blue) and juveniles (open circles). Sampling gaps are indicated with N letters, and sampling sessions without robins with blades. During the 14 years of sampling, 1376 different birds were captured (annual mean ± SD = 107.64 ± 32.11 individuals) although effective sample size was reduced due to occasionally missing morphological data.

**Figure S2.**

a

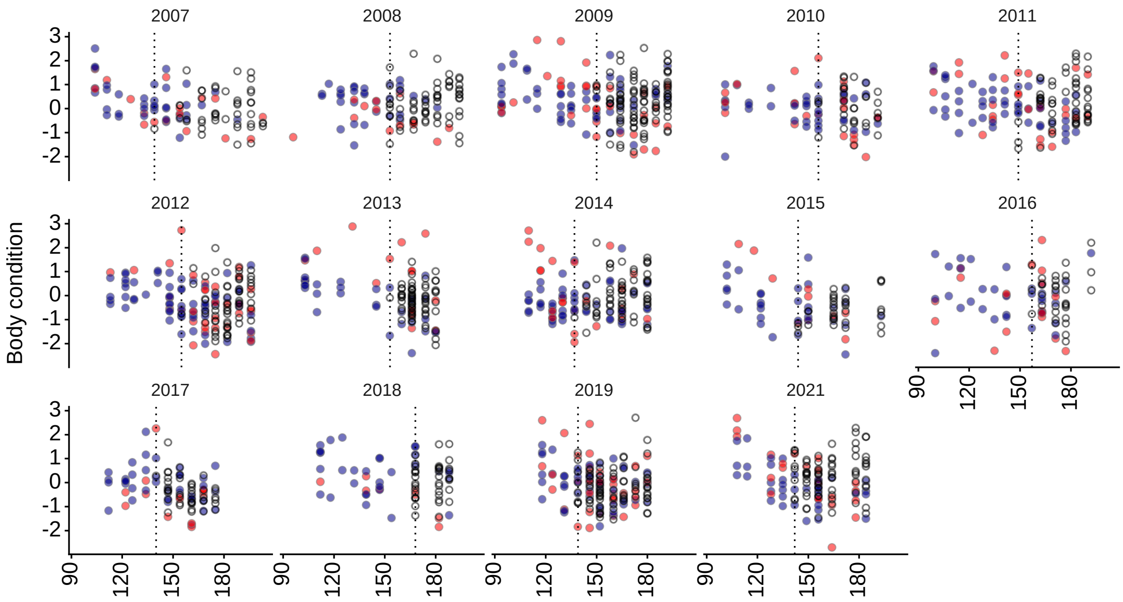
2007
2008
2009
2010
2011
2012
2013
2014
2015
2016
2017
2018
2019
2021
Body condition

b

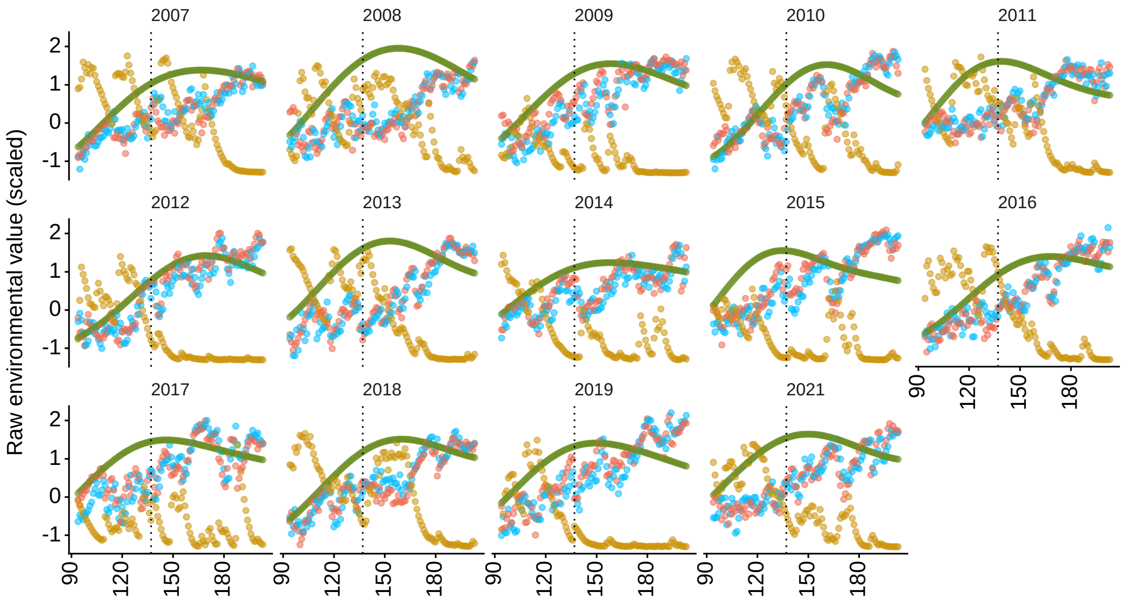
2007
2008
2009
2010
2011
2012
2013
2014
2015
2016
2017
2018
2019
2021
Raw environmental value (scaled)

c

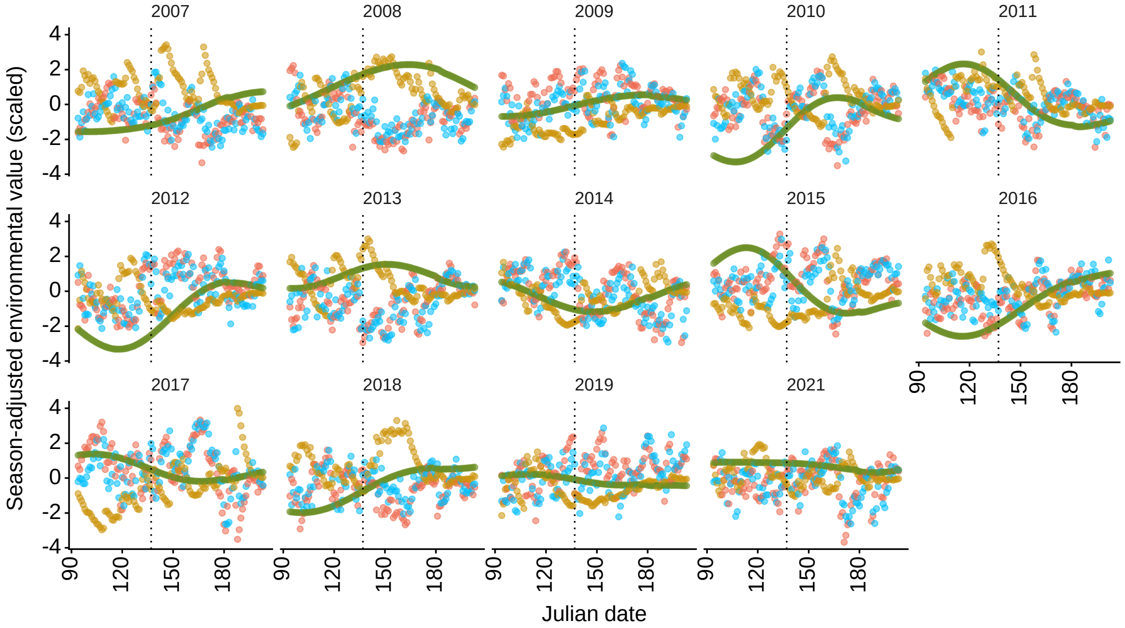


**Figure S2.** Seasonal variation in body condition and environmental variables each year. In body condition plots (a), individual values (dots) represent adult females (filled red), adult males (filled blue) and juveniles (filled grey). Vertical dashed lines represent the date of capture of the earliest juvenile each year, which divides each year in *pre-* and *post-fledging* seasons. Raw (b) and season-adjusted (c) values of the environmental variables used as potential correlates of body condition are shown with different colour fill: minimum temperature in blue, maximum temperature in red, soil moisture in ochre and EVI index in green. The vertical line common to all these plots represents the earliest capture date of a juvenile in any year, which sets the limit between *early* and *late* season all years.

### Text S1. Seasonal changes in within-individual body condition

We tested the significance of the within-individual change in body condition in each population group (adult males, adult females and juveniles) with repeated-measures mixed models using the *lme4* R package (Bates et al., 2015). Two analyses were performed, first considering all individual captures within a year (with any recapture interval), and second considering only short-time recaptures (maximum interval = 14 days). We included capture order (first or second) as a repeated-measures factor, and date of the last capture and recapture interval (in days) as covariates. The models included interactions between the covariates and recapture interval. Given that individuals with more than two captures in the same year had multiple recapture intervals, we controlled the non-independence of the data with a random factor that nested all alternative combinations of first and second capture within bird identity each year.

Within-year recapture interval in the whole population ranged 4-83 days for adults, and 6-38 days for juveniles. Considering all recaptures in the analyses, adult birds lost body condition between recaptures (mean ± SD of the first and second capture of alternative combinations; males = 0.26 ± 0.82 and -0.20 ± 0.85, females = 0.58 ± 1.14 and -0.21 ± 1.03, Table S1, Figure S3a). In adult males this loss was greater earlier in the season (estimate ± SE for date of last capture = 0.14 ± 0.05, Table S1) and with increasing recapture intervals (estimate ± SE = -0.30 ± 0.05, Table S1), while in the female body condition decreased more strongly towards the end of the season (estimate ± SE for date of last capture = -0.49 ± 0.10, Table S1). Juveniles slightly increased body condition between recaptures, more so with increasing recapture intervals (estimate ± SE = 0.24 ± 0.06, Table S1, Figure S3a). In addition, when we analysed short-time recaptures, we still detected individual body mass loss in adults but no change in juveniles (Table S1, Figure S3b). Nevertheless, this loss was consistent along the season and was only significantly influenced by recapture interval in adult males (Table S1).

**Table S1.** Results of repeated-measures mixed models assessing within-individual variation in body mass in relation to Julian date of the last capture and recapture interval (days elapsed between captures). A random factor (recapture time, REC) nested all alternative combinations of first and second capture of each individual each year within bird identity. Degrees of freedom were calculated using the Satterhwaite's method.

| | Adult males | Adult females | Juveniles |
|---|---|---|---|
| **All recaptures** | | | |
| Julian date of the last capture | $F_{1,592.0} = 70.20$; $p < 0.001$ | $F_{1,250.1} = 63.02$; $p < 0.001$ | $F_{1,125.3} = 1.29$; $p = 0.26$ |
| Recapture interval | $F_{1,580.1} = 23.72$; $p < 0.001$ | $F_{1,281.6} = 1.03$; $p = 0.31$ | $F_{1,200.9} = 0.002$; $p = 0.97$ |
| Recapture time (REC) | $F_{1,500.7} = 101.35$; $p < 0.001$ | $F_{1,250.3} = 45.54$; $p < 0.001$ | $F_{1,119.3} = 3.31$; $p = 0.07$ |
| Julian date of the last capture:REC | $F_{1,504.70} = 6.95$; $p < 0.01$ | $F_{1,252.2} = 23.65$; $p < 0.001$ | $F_{1,119.4} = 2.64$; $p = 0.11$ |
| Recapture interval:REC | $F_{1,505.4} = 29.96$; $p < 0.001$ | $F_{1,239.5} = 1.61$; $p = 0.21$ | $F_{1,119.5} = 14.04$; $p < 0.001$ |
| **Short-time recaptures (interval ≤ 14 days)** | | | |
| Julian date of the last capture | $F_{1,151.1} = 19.60$; $p < 0.001$ | $F_{1,48.8} = 36.39$; $p < 0.001$ | $F_{1,94.9} = 0.03$; $p = 0.87$ |
| Days between recaptures | $F_{1,158.8} = 0.40$; $p = 0.53$ | $F_{1,42.6} = 1.64$; $p = 0.21$ | $F_{1,130.7} = 0.03$; $p = 0.87$ |
| Recapture time (REC) | $F_{1,122.3} = 7.76$; $p < 0.01$ | $F_{1,50.7} = 8.75$; $p < 0.01$ | $F_{1,90.1} = 0.91$; $p = 0.34$ |
| Julian date of the last capture:REC | $F_{1,122.6} = 2.59$; $p = 0.11$ | $F_{1,55.1} = 0.88$; $p = 0.35$ | $F_{1,89.7} = 1.87$; $p = 0.18$ |

| | | | |
|---|---|---|---|
| Recapture interval:REC | $F_{1,122.1} = 4.23$; $p = 0.04$ | $F_{1,51.1} = 0.06$; $p = 0.81$ | $F_{1,89.8} = 0.14$; $p = 0.71$ |

**Figure S3.**

**a**

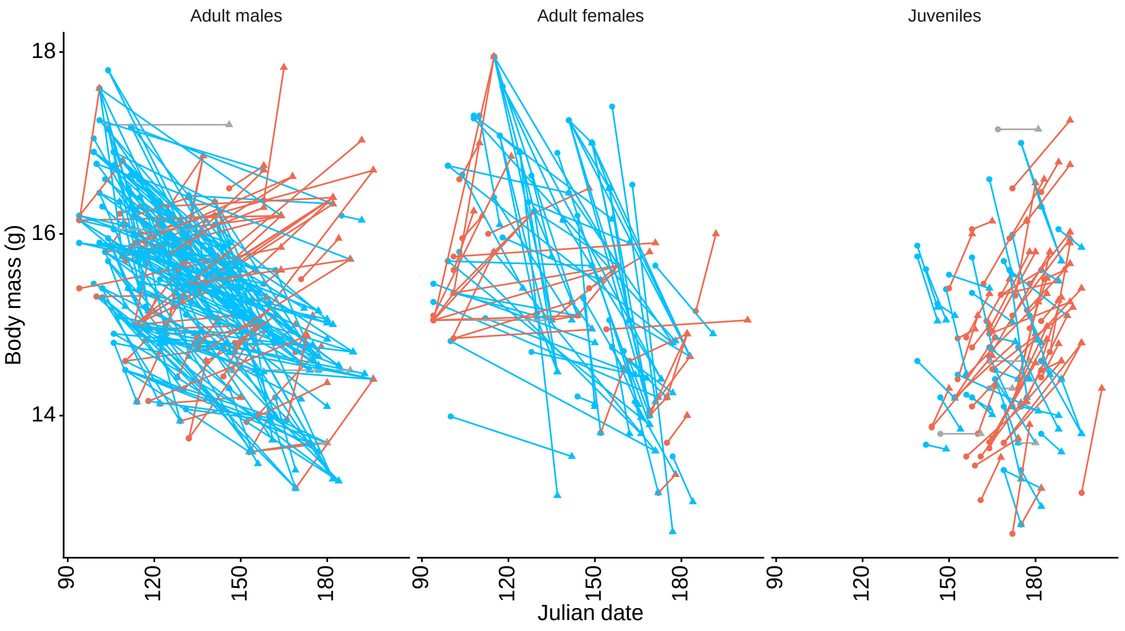


**b**

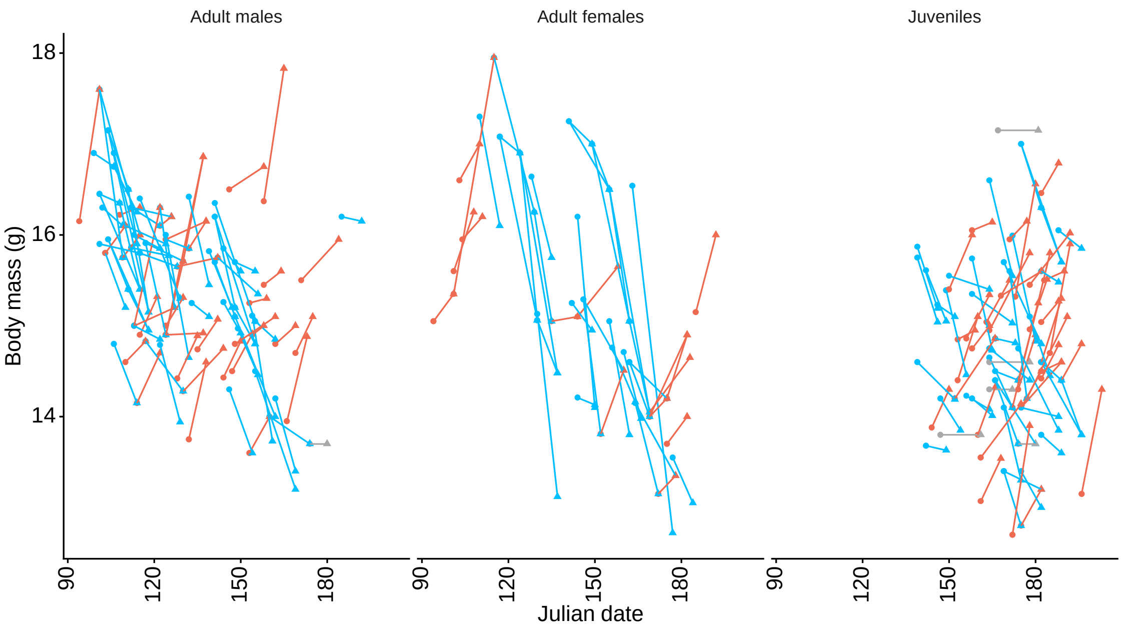


**Figure S3.** Trajectories of body mass change considering all (a) and short-time (b) individual recapture histories within one year for adult males, adult females and juveniles. The sign of the body condition change in each alternative combination is indicated with colours, blue for declining body mass, red for gain and grey for no change. We considered short-time lapse a maximum difference of 14 days between recaptures.

### CLIMWIN RESULTS:

**Table S2.** Identification of critical time windows (CTW) of season-adjusted environmental variables influencing body condition in adult male robins during the pre-fledging period, ranked according to their plausibility based on ΔAIC relative to the baseline model (ΔAICc for model $i = AICc_i - AICc_{baseline}$). The table shows the best model (the one with the lowest ΔAICc) for linear (lin) and quadratic (quad) relationships between the mean value of the environmental predictor during the CTW and body condition. Quadratic relationships are deemed more reliable than linear relationships only when they significantly improve model fit according to LRT tests. The location in time of the CTW is described by the days before capture date when the window opens and closes. Best models are highlighted in bold when significant based on $p_{\Delta AICc}$, resulting from the comparison of model ΔAICc with an expected distribution of ΔAICc values obtained from 100 random models with no relationship between weather and body condition. The percentage of models falling within the 95% confidence set based on cumulative AIC weights is shown only for best-supported environmental influences when significant. Environmental variables with ΔAIC > −2 relative to the baseline model were not further tested.

*Best models for body condition in **adult males** during the **pre-fledging period**:*

| Season-adjusted variable | Function | ΔAICc | CTW open | CTW close | *p* | % in 95CI |
|---|---|---|---|---|---|---|
| **Minimum temperature** | **quad** | **-21.91** | **13** | **10** | **<0.001** | **2%** |
| **Maximum temperature*** | **quad** | **-12.41** | **14** | **14** | **0.03** | |
| **Maximum temperature*** | **lin** | **-12.18** | **14** | **14** | **<0.001** | |
| Minimum temperature | lin | -5.84 | 11 | 10 | 0.07 | |
| Soil moisture | lin | -3.27 | 1 | 1 | 0.18 | |
| Soil moisture | quad | -2.68 | 1 | 1 | 0.31 | |
| EVI | lin | 0.39 | 25 | 25 | | |
| EVI | quad | 1.12 | 2 | 1 | | |

* The environmental predictor in this model was highly correlated with the top variable (r ≥ 0.66)

**Fig S4.** Plots describing the results of the best model (the one with the lowest ΔAICc) for the most reliable environmental influence on the body condition in adult male robins during the pre-fledging period (the quadratic relationship with season-adjusted minimum temperature). The figure shows the outcome of the function plotall() of the *climwin* R package, from top left to bottom right: (a) ΔAICc values of models considering every possible critical time window (CTW) during the 30 days before capture when compared to the baseline model without the climatic variable. (b) Percentage of all models falling within the 95% confidence set based on cumulative AICc weights. Coefficients for every CTW of linear (c) and quadratic (d) relationships between season-adjusted minimum temperature and body condition. (e) ΔAICc value of the best model (dashed line) compared with a distribution of ΔAICc values obtained from 100 randomized datasets with no relationship between weather and body condition. (f) Mean window open and close days of the best models included in the 95% confidence set. (g) Most reliable relationship between body condition (biological response) and minimum temperature (climate variable). The best model supported a CTW spanning from day 13 to 10 before capture.

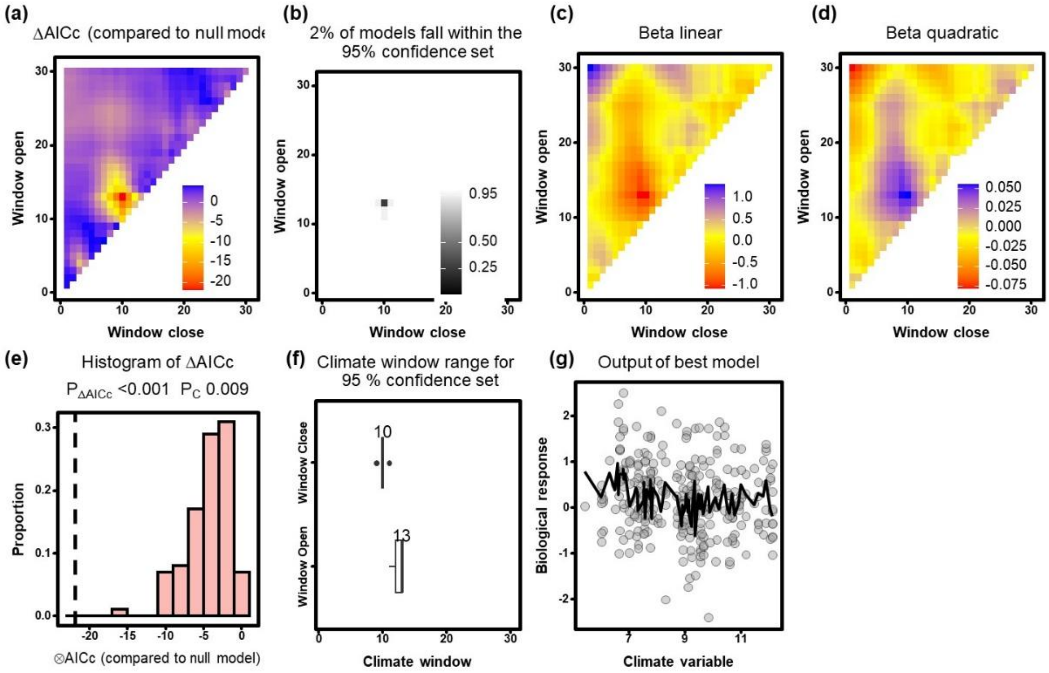

**Table S3.** Identification of critical time windows (CTW) of season-adjusted environmental variables influencing body condition in adult male robins during the post-fledging period, ranked according to their plausibility based on ΔAIC relative to the baseline model (ΔAICc for model $i$ = $AICc_i$ - $AICc_{baseline}$). The table shows the best model (the one with the lowest ΔAICc) for linear (lin) and quadratic (quad) relationships between the mean value of the environmental predictor during the CTW and body condition. Quadratic relationships are deemed more reliable than linear relationships only when they significantly improve model fit according to LRT tests. The location in time of the CTW is described by the days before capture date when the window opens and closes. Best models are highlighted in bold when significant based on $p_{\Delta AICc}$, resulting from the comparison of model ΔAICc with an expected distribution of ΔAICc values obtained from 100 random models with no relationship between weather and body condition. The percentage of models falling within the 95% confidence set based on cumulative AIC weights is shown only for best-supported environmental influences when significant. Environmental variables with ΔAIC > −2 relative to the baseline model were not further tested.

*Best models for body condition in* ***adult males*** *during the* ***post-fledging period****:*

| Season-adjusted variable | Function | ΔAICc | CTW open | CTW close | *p* | % in 95CI |
|---|---|---|---|---|---|---|
| **Soil moisture*** | **quad** | **-6.76** | **23** | **1** | **0.05** | **68%** |
| **Soil moisture*** | **lin** | **-6.23** | **23** | **1** | **<0.001** | **73%** |
| Minimum temperature | lin | -4.15 | 26 | 26 | 0.24 | |
| Maximum temperatura | lin | -3.99 | 29 | 29 | 0.13 | |
| Maximum temperatura | quad | -3.68 | 29 | 29 | 0.42 | |
| Minimum temperature | quad | -2.18 | 26 | 26 | 0.59 | |
| EVI | lin | -1.54 | 6 | 6 | | |
| EVI | quad | 0.22 | 12 | 4 | | |

* A quadratic relationship did not significantly improve model fit (LRT = 2.63, $p$ = 0.11), so the linear relationship was selected as most reliable.

**Fig S5.** Plots describing the results of the best model (the one with the lowest ΔAICc) for the most reliable environmental influence on the body condition in adult male robins during the post-fledging period (the linear relationship with season-adjusted soil moisture). The figure shows the outcome of the function plotall() of the *climwin* R package, from top left to bottom right: (a) ΔAICc values of models considering every possible critical time window (CTW) during the 30 days before capture when compared to the baseline model without the climatic variable. (b) Percentage of all models falling within the 95% confidence set based on cumulative AICc weights. Coefficients for every CTW of linear (c) and quadratic (d) relationships between season-adjusted soil moisture and body condition. (e) ΔAICc value of the best model (dashed line) compared with a distribution of ΔAICc values obtained from 100 randomized datasets with no relationship between weather and body condition. (f) Mean window open and close days of the best models included in the 95% confidence set. (g) Most reliable relationship between body condition (biological response) and soil moisture (climate variable). The best model supported a CTW spanning from day 23 to 1 before capture, although average CTW open and close days were 21 and 8, respectively.

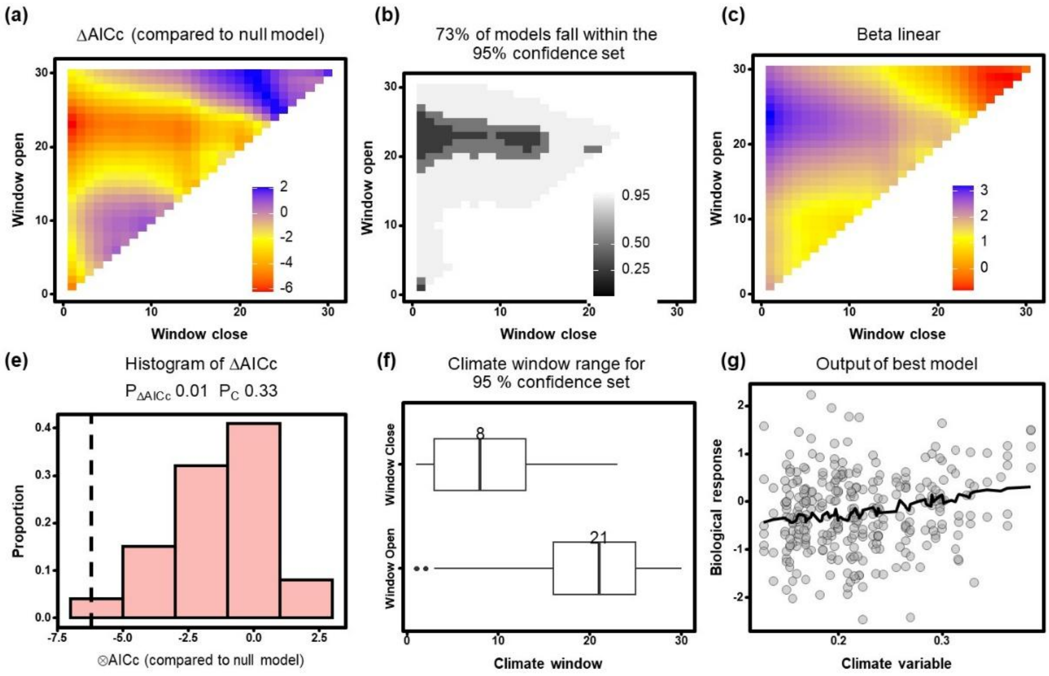

**Table S4.** Identification of critical time windows (CTW) of season-adjusted environmental variables influencing body condition in adult female robins during the pre-fledging period, ranked according to their plausibility based on ΔAIC relative to the baseline model (ΔAICc for model $i$ = $AICc_i$ - $AICc_{baseline}$). The table shows the best model (the one with the lowest ΔAICc) for linear (lin) and quadratic (quad) relationships between the mean value of the environmental predictor during the CTW and body condition. Quadratic relationships are deemed more reliable than linear relationships only when they significantly improve model fit according to LRT tests. The location in time of the CTW is described by the days before capture date when the window opens and closes. Best models are highlighted in bold when significant based on $p_{\Delta AICc}$, resulting from the comparison of model ΔAICc with an expected distribution of ΔAICc values obtained from 100 random models with no relationship between weather and body condition. The percentage of models falling within the 95% confidence set based on cumulative AIC weights is shown only for best-supported environmental influences when significant. Environmental variables with ΔAIC > −2 relative to the baseline model were not further tested.

*Best models for body condition in* ***adult females*** *during the* ***pre-fledging period****:*

| Season-adjusted variable | Function | ΔAICc | CTW open | CTW close | $p$ | % in 95CI |
|---|---|---|---|---|---|---|
| **Soil moisture** | **quad** | **-8.77** | **7** | **1** | **0.01** | **40%** |
| Minimum temperature | quad | -7.76 | 22 | 2 | 0.20 | |
| Minimum temperature | lin | -6.44 | 3 | 3 | 0.14 | |
| Maximum temperature | quad | -4.80 | 30 | 8 | 0.35 | |
| Maximum temperature | lin | -3.69 | 27 | 18 | 0.27 | |
| **EVI** | **lin** | **-2.54** | **1** | **1** | **0.03** | **89%** |
| EVI | quad | -0.24 | 1 | 1 | | |
| Soil moisture | lin | 0.06 | 30 | 30 | | |

**Fig S6.** Plots describing the results of the best model (the one with the lowest ΔAICc) for the most reliable environmental influence on the body condition in adult female robins during the pre-fledging period (the quadratic relationship with season-adjusted soil moisture). The figure shows the outcome of the function plotall() of the *climwin* R package, from top left to bottom right: (a) ΔAICc values of models considering every possible critical time window (CTW) during the 30 days before capture when compared to the baseline model without the climatic variable. (b) Percentage of all models falling within the 95% confidence set based on cumulative AICc weights. Coefficients for every CTW of linear (c) and quadratic (d) relationships between season-adjusted soil moisture and body condition. (e) ΔAICc value of the best model (dashed line) compared with a distribution of ΔAICc values obtained from 100 randomized datasets with no relationship between weather and body condition. (f) Mean window open and close days of the best models included in the 95% confidence set. (g) Most reliable relationship between body condition (biological response) and soil moisture (climate variable). The best model supported a CTW spanning from day 7 to 1 before capture, although average CTW open and close days were 15 and 5, respectively.

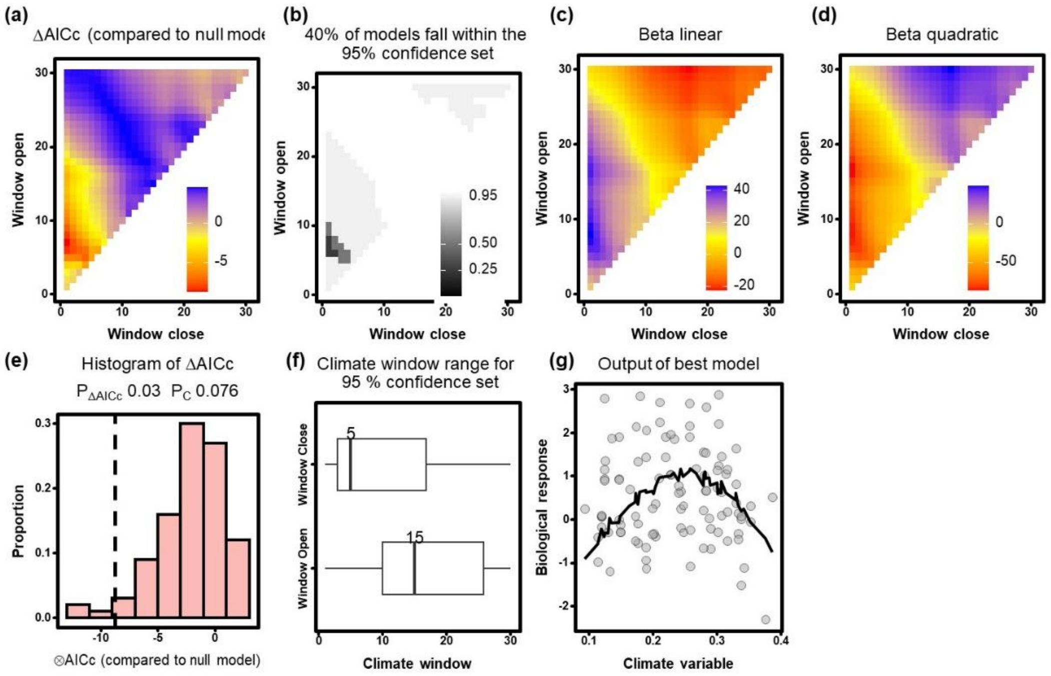

**Table S5.** Identification of critical time windows (CTW) of season-adjusted environmental variables influencing body condition in adult female robins during the post-fledging period, ranked according to their plausibility based on ΔAIC relative to the baseline model (ΔAICc for model $i$ = $AICc_i$ - $AICc_{baseline}$). The table shows the best model (the one with the lowest ΔAICc) for linear (lin) and quadratic (quad) relationships between the mean value of the environmental predictor during the CTW and body condition. Quadratic relationships are deemed more reliable than linear relationships only when they significantly improve model fit according to LRT tests. The location in time of the CTW is described by the days before capture date when the window opens and closes. Best models are highlighted in bold when significant based on $p_{\Delta AICc}$, resulting from the comparison of model ΔAICc with an expected distribution of ΔAICc values obtained from 100 random models with no relationship between weather and body condition. The percentage of models falling within the 95% confidence set based on cumulative AIC weights is shown only for best-supported environmental influences when significant. Environmental variables with ΔAIC > −2 relative to the baseline model were not further tested.

*Best models for body condition in* ***adult females*** *during the* ***post-fledging period****:*

| Season-adjusted variable | Function | ΔAICc | CTW open | CTW close | *p* | % in 95CI* |
|---|---|---|---|---|---|---|
| Minimum temperature | lin | -4.87 | 13 | 13 | 0.19 | |
| Maximum temperature | lin | -4.67 | 24 | 24 | 0.16 | |
| Soil moisture | lin | -4.45 | 24 | 24 | 0.08 | |
| Maximum temperature | quad | -4.29 | 18 | 14 | 0.29 | |
| Soil moisture | quad | -3.23 | 24 | 24 | 0.25 | |
| Minimum temperature | quad | -2.89 | 25 | 12 | 0.59 | |
| EVI | quad | -0.25 | 16 | 16 | | |
| EVI | lin | 1.60 | 30 | 30 | | |

* None of the models was significant.

**Table S6.** Identification of critical time windows (CTW) of season-adjusted environmental variables influencing body condition in juvenile robins, ranked according to their plausibility based on ΔAIC relative to the baseline model (ΔAICc for model $i$ = $AICc_i$ - $AICc_{baseline}$). The table shows the best model (the one with the lowest ΔAICc) for linear (lin) and quadratic (quad) relationships between the mean value of the environmental predictor during the CTW and body condition. Quadratic relationships are deemed more reliable than linear relationships only when they significantly improve model fit according to LRT tests. The location in time of the CTW is described by the days before capture date when the window opens and closes. Best models are highlighted in bold when significant based on $p_{\Delta AICc}$, resulting from the comparison of model ΔAICc with an expected distribution of ΔAICc values obtained from 100 random models with no relationship between weather and body condition. The percentage of models falling within the 95% confidence set based on cumulative AIC weights is shown only for best-supported environmental influences when significant. Environmental variables with ΔAIC > −2 relative to the baseline model were not further tested.

*Best models for body condition in* ***juveniles****:*

| Season-adjusted variable | Function | ΔAICc | CTW open | CTW close | *p* | % in 95CI |
|---|---|---|---|---|---|---|
| **Minimum temperature** | **lin** | **-8.37** | **6** | **2** | **0.01** | **40%** |
| **Minimum temperature*** | **quad** | **-8.19** | **5** | **5** | **0.06** | |
| **Maximum temperature*** | **lin** | **-6.43** | **9** | **1** | **0.05** | |
| Maximum temperature | quad | -4.55 | 6 | 6 | 0.12 | |
| Soil moisture | quad | -3.50 | 15 | 5 | 0.11 | |
| Soil moisture | lin | -2.46 | 7 | 7 | 0.14 | |
| EVI | lin | 1.40 | 10 | 10 | | |
| EVI | quad | 2.05 | 13 | 1 | | |

* The environmental predictor in this model was highly correlated with the top variable ($r \geq 0.88$)

**Fig S7.** Plots describing the results of the best model (the one with the lowest ΔAICc) for the most reliable environmental influence on the body condition in juvenile robins (the linear relationship with season-adjusted minimum temperature). The figure shows the outcome of the function plotall() of the *climwin* R package, from top left to bottom right: (a) ΔAICc values of models considering every possible critical time window (CTW) during the 30 days before capture when compared to the baseline model without the climatic variable. (b) Percentage of all models falling within the 95% confidence set based on cumulative AICc weights. Coefficients for every CTW of linear (c) and quadratic (d) relationships between season-adjusted minimum temperature and body condition. (e) ΔAICc value of the best model (dashed line) compared with a distribution of ΔAICc values obtained from 100 randomized datasets with no relationship between weather and body condition. (f) Mean window open and close days of the best models included in the 95% confidence set. (g) Most reliable relationship between body condition (biological response) and minimum temperature (climate variable). The best model supported a CTW spanning from day 6 to 2 before capture, although average CTW open and close days were 8 and 2, respectively.

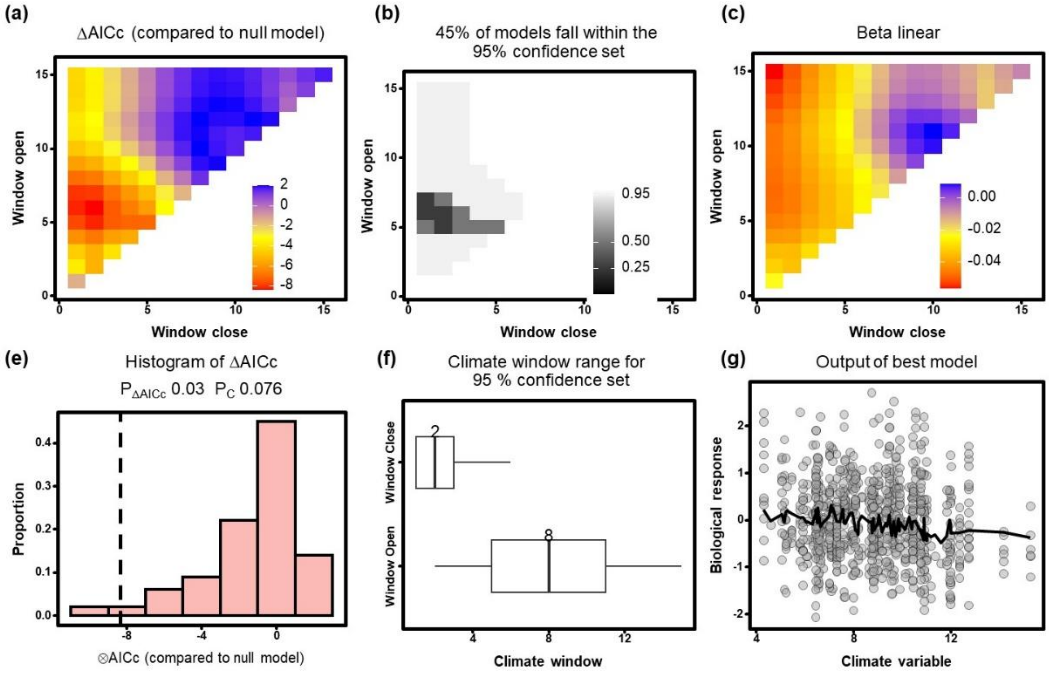

**Table S7.** Identification of critical time windows (CTW) of season-adjusted environmental variables influencing within-individual variation in body condition in adult male robins, ranked according to their plausibility based on ΔAIC relative to the baseline model (ΔAICc for model $i = AICc_i - AICc_{baseline}$). The table shows the best model (the one with the lowest ΔAICc) for linear (lin) and quadratic (quad) relationships between the mean value of the environmental predictor during the CTW and body condition. Quadratic relationships are deemed more reliable than linear relationships only when they significantly improve model fit according to LRT tests. The location in time of the CTW is described by the days before capture date when the window opens and closes. Best models are highlighted in bold when significant based on $p_{\Delta AICc}$, resulting from the comparison of model ΔAICc with an expected distribution of ΔAICc values obtained from 100 random models with no relationship between weather and body condition. The percentage of models falling within the 95% confidence set based on cumulative AIC weights is shown only for best-supported environmental influences when significant. Environmental variables with ΔAIC > −2 relative to the baseline model were not further tested.

*Best models for* ***within-individual*** *change in body condition in* ***adult males****:*

| Season-adjusted variable | Function | ΔAICc | CTW open | CTW close | *p* | % in 95CI* |
|---|---|---|---|---|---|---|
| Maximum temperature | quad | -7.14 | 7 | 7 | 0.08 | |
| Minimum temperature | lin | -4.24 | 14 | 14 | 0.16 | |
| Minimum temperature | quad | -2.50 | 14 | 13 | 0.42 | |
| Soil moisture | quad | -1.05 | 6 | 6 | | |
| Maximum temperature | lin | 0.62 | 14 | 14 | | |
| Soil moisture | lin | 1.32 | 8 | 8 | | |
| EVI | lin | 2.27 | 14 | 14 | | |
| EVI | quad | 3.76 | 1 | 1 | | |

* None of the models was significant.

**Table S8.** Identification of critical time windows (CTW) of season-adjusted environmental variables influencing within-individual variation in body condition in adult female robins, ranked according to their plausibility based on ΔAIC relative to the baseline model (ΔAICc for model $i$ = $AICc_i$ - $AICc_{baseline}$). The table shows the best model (the one with the lowest ΔAICc) for linear (lin) and quadratic (quad) relationships between the mean value of the environmental predictor during the CTW and body condition. Quadratic relationships are deemed more reliable than linear relationships only when they significantly improve model fit according to LRT tests. The location in time of the CTW is described by the days before capture date when the window opens and closes. Best models are highlighted in bold when significant based on $p_{\Delta AICc}$, resulting from the comparison of model ΔAICc with an expected distribution of ΔAICc values obtained from 100 random models with no relationship between weather and body condition. The percentage of models falling within the 95% confidence set based on cumulative AIC weights is shown only for best-supported environmental influences when significant. Environmental variables with ΔAIC > −2 relative to the baseline model were not further tested.

*Best models for **within-individual** change in body condition in **adult females**:*

| Season-adjusted variable | Function | ΔAICc | CTW open | CTW close | *p* | % in 95CI* |
|---|---|---|---|---|---|---|
| Maximum temperature | quad | -5.77 | 11 | 9 | 0.08 | |
| Minimum temperature | quad | -1.93 | 5 | 4 | | |
| Soil moisture | quad | -1.88 | 10 | 10 | | |
| Maximum temperature | lin | 0.02 | 7 | 5 | | |
| Minimum temperature | lin | 1.41 | 6 | 4 | | |
| Soil moisture | lin | 1.89 | 1 | 1 | | |
| EVI | quad | 2.52 | 11 | 11 | | |
| EVI | lin | 2.53 | 1 | 1 | | |

* None of the models were significant.

**Table S9.** Identification of critical time windows (CTW) of season-adjusted environmental variables influencing within-individual variation in body condition in juvenile robins, ranked according to their plausibility based on ΔAIC relative to the baseline model (ΔAICc for model $i$ = $AICc_i$ - $AICc_{baseline}$). The table shows the best model (the one with the lowest ΔAICc) for linear (lin) and quadratic (quad) relationships between the mean value of the environmental predictor during the CTW and body condition. Quadratic relationships are deemed more reliable than linear relationships only when they significantly improve model fit according to LRT tests. The location in time of the CTW is described by the days before capture date when the window opens and closes. Best models are highlighted in bold when significant based on $p_{\Delta AICc}$, resulting from the comparison of model ΔAICc with an expected distribution of ΔAICc values obtained from 100 random models with no relationship between weather and body condition. The percentage of models falling within the 95% confidence set based on cumulative AIC weights is shown only for best-supported environmental influences when significant. Environmental variables with ΔAIC > −2 relative to the baseline model were not further tested.

*Best models for* ***within-individual*** *change in body condition in* ***juveniles****:*

| Season-adjusted variable | Function | ΔAICc | CTW open | CTW close | *p* | % in 95CI* |
|---|---|---|---|---|---|---|
| Minimum temperature | lin | -2.24 | 11 | 11 | 0.18 | |
| Minimum temperature | quad | -1.77 | 11 | 11 | | |
| Maximum temperature | lin | -1.54 | 5 | 3 | | |
| Soil moisture | quad | -1.04 | 3 | 2 | | |
| Soil moisture | lin | -0.92 | 3 | 2 | | |
| Maximum temperature | quad | 0.45 | 5 | 4 | | |
| EVI | lin | 2.16 | 14 | 14 | | |
| EVI | quad | 4.25 | 14 | 14 | | |

* None of the models were significant.

**Table S10.** Results of the models analysing linear long-term trends of body condition in adult males, adult females and juveniles, without (*basic models*) and with environmental variables selected in weather sliding-window analyses (*full environmental models*). Data were fitted with generalized mixed models (*mgcv* package; Wood, 2017) and included the linear effect of year and Julian date of capture as a basic model. Non-linear patterns were tested (F tests) only when quadratic effects (edf = 2) of environmental variables were deemed most reliable. Year, Julian date and environmental variables were previously standardized. Different models were fit for adults using data from the pre-fledging (PRE) and post-fledging periods (POST). The sample size of each model is indicated (n), and dbc stands for days before capture of critical time windows for environmental influences.

| | edf | $R^2_{adj}$ | Estimate | SE | t | F | *p* |
|---|---|---|---|---|---|---|---|
| Basic models | | | | | | | |
| *Between-individuals models* | | | | | | | |
| Adult males-PRE (n = 287) | | 0.072 | | | | | |
| Year | | | -0.007 | 0.054 | -0.13 | | 0.89 |
| Julian date | | | -0.254 | 0.036 | -7.01 | | <0.001 |
| Adult males-POST (n = 303) | | 0.026 | | | | | |
| Year | | | -0.156 | 0.049 | -3.16 | | 0.002 |
| Julian date | | | -0.039 | 0.041 | -0.96 | | 0.34 |
| Adult females-PRE (n = 100) | | 0.023 | | | | | |
| Year | | | 0.020 | 0.116 | 0.18 | | 0.86 |
| Julian date | | | -0.205 | 0.102 | -2.02 | | 0.047 |
| Adult females-POST (n = 194) | | 0.025 | | | | | |
| Year | | | -0.082 | 0.082 | -1.00 | | 0.32 |
| Julian date | | | -0.227 | 0.079 | -2.87 | | 0.005 |
| Juveniles (n = 810) | | 0.015 | | | | | |
| Year | | | -0.008 | 0.033 | -0.26 | | 0.80 |
| Julian date | | | 0.122 | 0.030 | 4.01 | | <0.001 |
| *Within-individuals models* | | | | | | | |
| Adult male (n = 90) | | 0.037 | | | | | |
| Year | | | 0.143 | 0.078 | 1.82 | | 0.07 |
| Julian date | | | 0.091 | 0.078 | 1.17 | | 0.25 |
| Adult female (n = 46) | | -0.008 | | | | | |
| Year | | | -0.195 | 0.177 | -1.10 | | 0.28 |
| Julian date | | | -0.064 | 0.176 | -0.37 | | 0.72 |
| Juveniles (n = 79) | | -0.006 | | | | | |
| Year | | | 0.024 | 0.080 | 0.30 | | 0.76 |
| Julian date | | | 0.124 | 0.080 | 1.55 | | 0.13 |
| Full environmental models | | | | | | | |
| *Between-individuals models* | | | | | | | |
| Adult males-PRE (n = 287) | | 0.104 | | | | | |
| Year | | | 0.024 | 0.054 | 0.45 | | 0.66 |
| Julian date | | | -0.279 | 0.033 | -8.35 | | <0.001 |
| Minimum temperature (13-10 dbc) | 2 | | | | | 14.85 | <0.001 |
| Adult males-POST (n = 303) | | 0.041 | | | | | |

| | | | | | | | |
|---|---|---|---|---|---|---|---|
| Year | | | -0.111 | 0.052 | -2.16 | | 0.03 |
| Julian date | | | -0.049 | 0.040 | -1.22 | | 0.22 |
| Minimum temperature (13-10 dbc) | | | 0.124 | 0.046 | 2.72 | | 0.007 |
| Adult females-PRE (n = 100) | | 0.074 | | | | | |
| Year | | | -0.065 | 0.118 | -0.56 | | 0.58 |
| Julian date | | | -0.163 | 0.101 | -1.62 | | 0.11 |
| Soil moisture (7-1 dbc) | 2 | | | | | 3.9 | 0.02 |
| Adult females-POST * | | | | | | | |
| Juveniles (n = 810) | | 0.027 | | | | | |
| Year | | | 0.007 | 0.033 | 0.21 | | 0.83 |
| Julian date | | | 0.122 | 0.030 | 4.04 | | <0.001 |
| Minimum temperature (6-2 dbc) | | | -0.097 | 0.027 | -3.63 | | <0.001 |
| *Within-individuals models* | | | | | | | |
| Adult male* | | | | | | | |
| Adult female* | | | | | | | |
| Juveniles* | | | | | | | |

* No full environmental model reported because no reliable climatic windows were selected for any environmental variable.

**Figure S8.** Long-term trends of season-adjusted (sadj) environmental variables (maximum temperature, minimum temperature, soil moisture and EVI) during the *early* (red) and *late* (ochre) breeding seasons across the period 2007–2021.

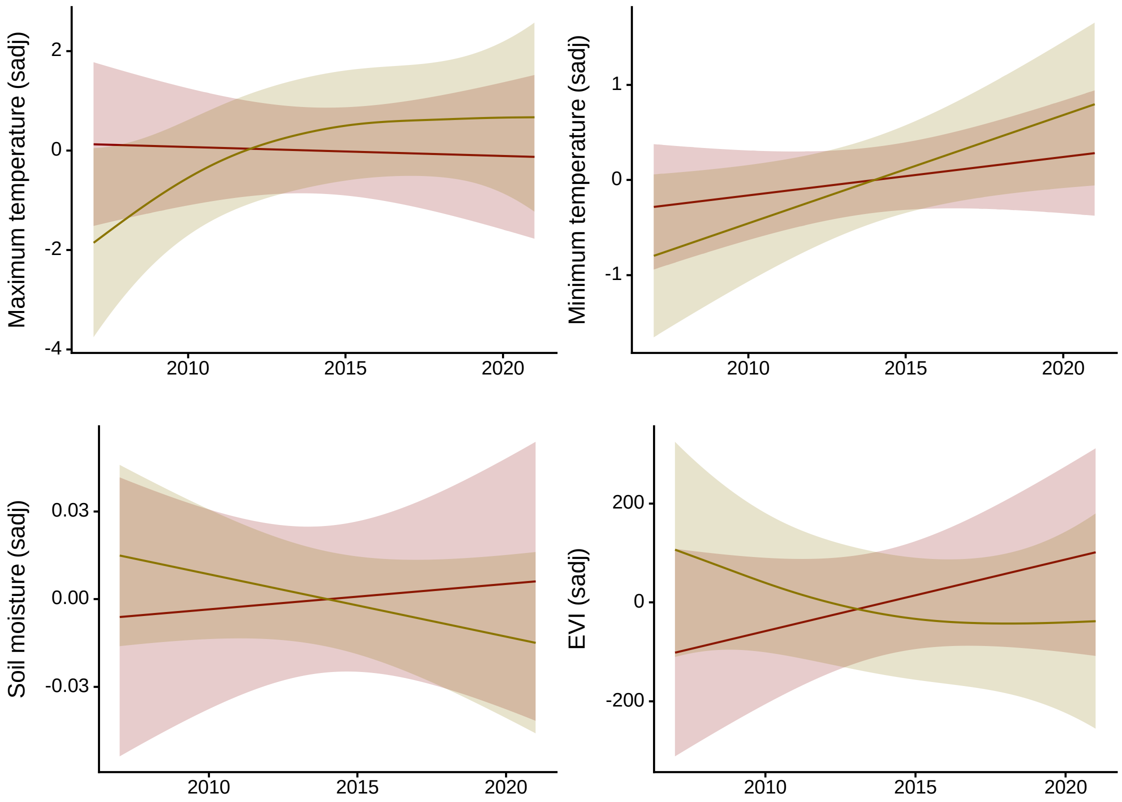

**Table S11**. Best Cormack-Jolly-Seber models (those with ΔAICc < 2) of variation in apparent survival (Φ) and probability of recapture (p) of adult and juvenile robins. We also show the best model that did not include body condition (the variable of interest) and the null model ($\Phi_{\text{intercept}} + p_{\text{intercept}}$). The AIC value, number of parameters (*k*) and Akaike weight are shown for each model.

| Model | *k* | AICc | ΔAICc | *weight* |
|---|---|---|---|---|
| Adults | | | | |
| 1: $\Phi_{\text{sex}} + p_{\text{intercept}}$ | 3 | 583.77 | 0.000 | 0.38 |
| 2: $\Phi_{\text{sex + adult body condition}} + p_{\text{intercept}}$ | 4 | 585.61 | 1.839 | 0.15 |
| 3: $\Phi_{\text{sex + Time}} + p_{\text{intercept}}$ | 4 | 585.69 | 1.918 | 0.15 |
| 4: $\Phi_{\text{intercept}} + p_{\text{intercept}}$ | 2 | 586.51 | 2.738 | 0.10 |
| Juveniles | | | | |
| 1: $\Phi_{\text{juvenile body condition}} + p_{\text{intercept}}$ | 3 | 346.66 | 0.000 | 0.41 |
| 2: $\Phi_{\text{juvenile body condition + Time}} + p_{\text{intercept}}$ | 4 | 348.65 | 1.997 | 0.15 |
| 4: $\Phi_{\text{intercept}} + p_{\text{intercept}}$ | 2 | 349.66 | 3.006 | 0.09 |

**Text S2. Comparison of body condition between birds of each population group that were recaptured or not**

We assessed whether body condition measured at first capture differed between birds that were or were not recaptured. General linear models were fitted for adult males, adult females and juveniles, always controlling for year (as a factor) and Julian capture date. For adults of either sex, we fitted different models using data from pre-fledging and post-fledging periods.

In juvenile robins, individuals that were recaptured had a higher body condition at first capture (mean ± SD = 0.19 ± 0.96) than those not seen again (-0.03 ± 0.81; $F_{1,702} = 5.77$, $p = 0.02$), controlling for an increase in body condition during the season (Julian date estimate ± SE = 0.12 ± 0.03, $F_{1,702} = 11.28$, $p < 0.001$) and significant differences in juvenile body condition between years ($F_{13,702} = 5.93$, $p < 0.001$). In adults, we did not detect significant differences in body condition between individuals that were recaptured or not (all comparisons with $p > 0.14$, which corresponded to adult males in the pre-fledging period), controlling for a decrease in body condition during the season (Julian date $p \leq 0.04$ for adult males in the pre-fledging period or adult females in the post-fledging period, other alternatives $p > 0.53$) and significant differences in adult males body condition between years ($p < 0.01$, females $p > 0.25$).
